# Different Sources of Expectations Differentially Modulate Communication Between Auditory and Prefrontal Cortices During Auditory Decision-Making

**DOI:** 10.64898/2026.07.22.740131

**Authors:** Corey Roach, Lalitta Suriya-Arunroj, Sonia Bilal, Sophia Fu, Joshua I. Gold, Yale Cohen

## Abstract

Perceptual decisions can be influenced by expectations that arise from different sources. However, relatively little is known about how and where in the brain these different sources affect decision formation, particularly in the auditory system. To begin to address this question, we recorded local field potentials (LFPs) simultaneously from the auditory cortex (AC) and ventrolateral prefrontal cortex (vlPFC) while monkeys performed an auditory frequency-discrimination task. We manipulated their expectations by presenting either informative cues (LEDs that predicted the target frequency) or uninformative cues (tone sequences that did not predict the target frequency) prior to target onset. Both cues had similar effects on the monkeys’ accuracy and response times but induced distinct neurophysiological signatures. Coherence, a measure of undirected coupling, between AC and vlPFC increased from cue to target presentation on informative trials but decreased from cue to target presentation on uninformative trials. Phase-slope index (PSI), a measure of directed coupling, between AC and vlPFC was strongest when the target tone contradicted expectations, particularly for ambiguous stimulus conditions when expectations were most valuable. These expectation-dependent changes in the magnitude of functional connectivity also included changes in its spatial organization. Together, these findings provide new insights into how different expectations can dynamically reconfigure functional connectivity between cortical brain areas to support different task demands and behavioral outcomes.

## Introduction

The ventral auditory pathway plays a causal role in auditory perceptual decision-making (Cohen et al., 2016; Plakke & Romanski, 2014; Tsunada et al., 2019). This pathway starts in the core region of the auditory cortex (AC), continues in the belt and parabelt regions, then projects to the ventrolateral prefrontal cortex (vlPFC) (Hackett et al., 1999; Romanski et al., 1999a; Romanski, et al., 1999b). Each of these regions contains neurons whose activity patterns relate to the transformation of auditory information into a decision that guides behavior (Rauschecker & Tian, 2000; Romanski, et al., 1999b; Tian et al., 2001; Tsunada et al., 2011; Tsunada & Cohen, 2014). In addition to this feedforward processing, this pathway includes rich patterns of feedback connectivity that are thought to contribute to biases in sensory gain and feature selectivity, sharpening of acoustic representations under noisy conditions, and flexible reweighing of feedforward evidence depending on task demands (Gerbella et al., 2010; Plakke & Romanski, 2014; Rauschecker & Afsahi, 2023). However, exactly how the functional connectivity patterns between different regions of this pathway support both feedforward and feedback components of decision processing is not known.

To address this issue, we tested how functional connectivity between AC and vlPFC (likely mediated via feedforward and feedback pathways through the belt and parabelt regions) is modulated by different forms of expectations. We recorded local-field potentials (LFPs) simultaneously in the AC and vlPFC of monkeys performing an auditory frequency-discrimination task. The monkeys reported whether a target tone embedded in a noisy background was low or high frequency. We manipulated task difficulty by varying the signal-to-noise ratio (SNR) between the target tone and the noisy background. We manipulated expectations in two ways. On informative trials, an LED color signaled the probability of the frequency of the target tone. On uninformative trials, a neutral LED followed by a sequence of tone bursts (“pretones”) were presented that did not predict the target-tone frequency.

Like for human participants performing a similar task (Tardiff et al., 2022), both the informative and uninformative cues biased choices and response times (RTs). However, these behavioral biases corresponded to different patterns of AC-vlPFC connectivity for the different sources of expectations. On informative trials, AC-vlPFC coherence was lower during presentation of the LED and higher during presentation of the target tone. In contrast, on uninformative trials, the pattern reversed, with higher coherence during presentation of the LED and lower coherence during presentation of the target tone. Furthermore, AC-vlPFC connectivity was modulated by congruence, including stronger coupling when the monkeys made choices that contradicted their initial expectations (e.g., choosing “low frequency” despite being cued to expect a high-frequency tone). Together, these findings illustrate how expectations can dynamically shape functional connectivity in the ventral auditory pathway to support auditory-driven behavior under varying task demands.

## Results

We trained two adult male monkeys to perform an auditory frequency-discrimination task with informative and uninformative cues (**Fig. 1**). Each trial began with a LED. On informative trials, a blue or green LED indicated, with 75% reliability, that the target tone was high or low frequency, respectively. On uninformative trials, a yellow LED was followed by 3 pretones, none of which were predictive of the target frequency. At any time after onset of the target tone, the monkeys could use the joystick to report their decision about whether the target tone was high or low frequency. They were rewarded for correct choices.

**Figure 1:**
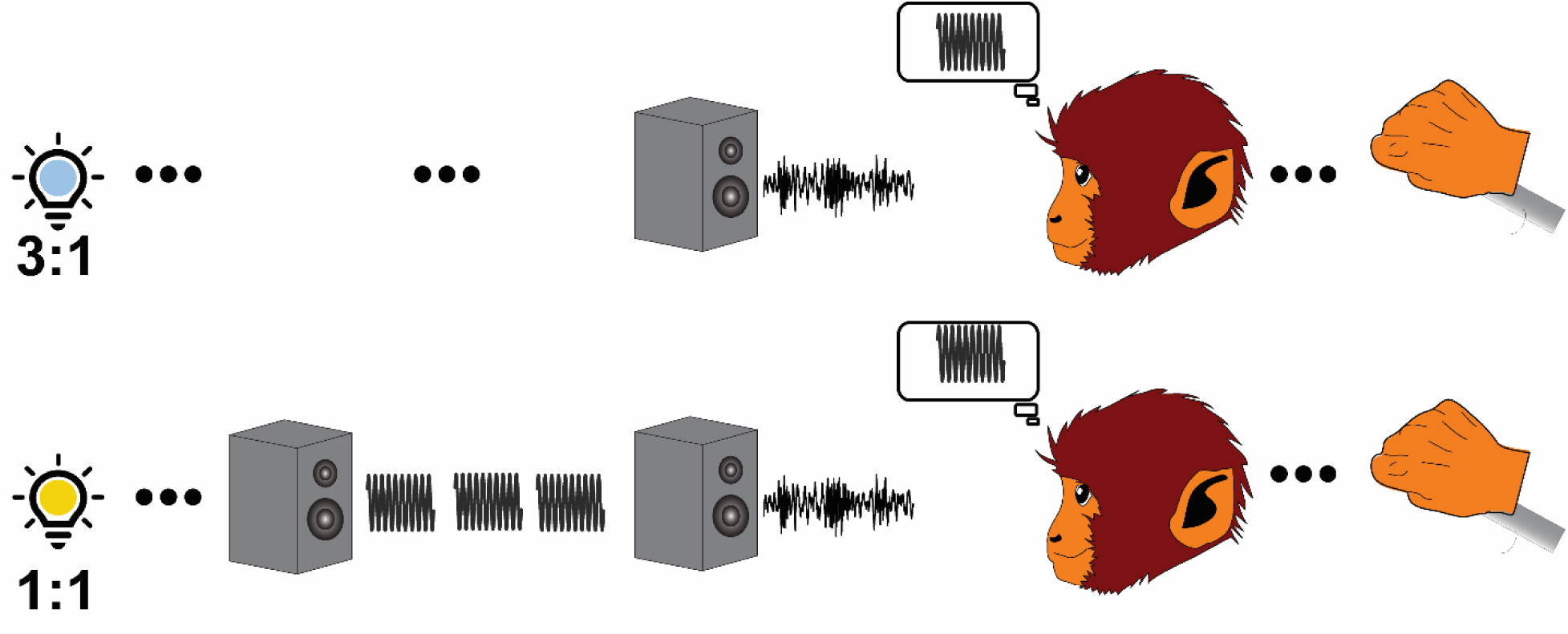
Schematic of the auditory frequency-discrimination task. Using a joystick, monkeys reported whether a target tone burst embedded in various amounts of background noise was “low frequency” (L) or “high frequency” (H). Prior to onset of the target tone, we presented informative or uninformative cues. On informative trials, an LED predicted the frequency of the target tone with 75% accuracy (H:L= 3:1 for blue, 1:3 for green). On uninformative trials, a “neutral” yellow LED (H:L= 1:1) was followed by a sequence of three tone bursts (“pretones”). The pretones were either all low frequency or high frequency and, like the neutral yellow LED, did not predict the frequency of the tone burst.

### Informative and uninformative cues bias behavioral choice and response time

The monkey’s decisions depended systematically on the SNR of the target tone, relative to the background noise. In general, on trials with (absolute value) high-SNR stimuli, the monkeys almost always reported the correct answer. As absolute SNR decreased, performance accuracy decreased. Accordingly, across all conditions, psychometric functions show increasing proportions of “high-frequency” choices as signed SNR increased (**Fig. 2**; **Table 1**). Response times (RTs) were also systematically modulated by SNR with the shortest RTs on trials with (absolute value) high-SNR values and longer RTs at lower SNR values, particularly on informative trials (**Fig. 2**, **Table 2**).

**Figure 2:**
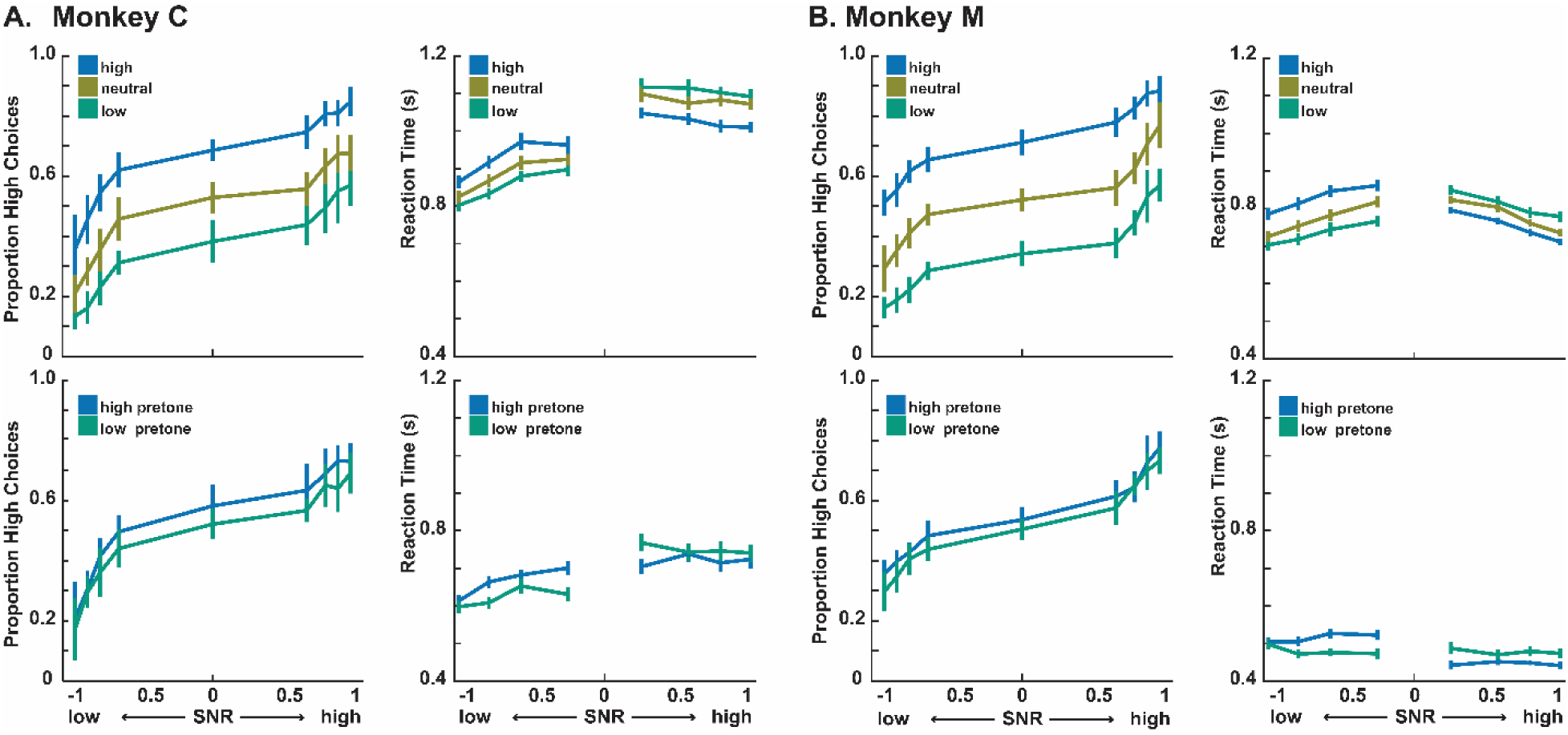
Psychometric and chronometric performance. For Monkey C (**A**) and Monkey M (**B**), the proportion of trials in which the monkey chose “high frequency” (psychometric functions, **left**) and response time (RT; chronometric function, **right**) are each plotted as a function of the signal-to-noise ratio (SNR) between the target tone and the background noise. Positive and negative SNRs indicate high- and low-frequency targets, respectively. 0 SNR indicates trials in which we presented only the background noise. **Top row**, Low- and high-frequency informative trials and uninformative trials (“neutral,” corresponding to all pretone frequencies; see legend). **Bottom row**, Uninformative trials with different frequency pretones (see legend). Lines are not fits but simply connect the data points for visualization. SNR values are scaled between 0 and 1. Negative or positive SNR values indicate that the target frequency was low frequency or high frequency, respectively. Error bars represent standard errors of the mean across sessions.

The monkeys’ choices and RTs also depended systematically on expectations provided by the informative and uninformative cues (**Fig. 2**). Specifically, during informative trials, the monkeys were more likely to choose “low frequency” (“high frequency”) when the LED cued a low-frequency (high-frequency) target as being more likely (**Table 3**). These choice effects were accompanied by RT effects that depended on the congruency between the informative cue and the target frequency. RTs were faster on congruent trials (e.g., trials in which the cue correctly signaled that the target was high frequency) than on incongruent trials (e.g., trials in which the LED signaled high frequency but the target was low frequency; Monkey C, congruent median=935 ms, incongruent median=1011 ms; Monkey M, congruent median =732 ms, incongruent median=813 ms), with RTs on uninformative trials intermediate between the two (**Table 4**).

The uninformative pretones resulted in similar, albeit smaller in magnitude, effects on both choice and RTs. The monkeys’ choices tended to be biased towards the frequency of the pretones (**Table 3**). These choice effects were accompanied by RT effects that also depended on congruency, including faster RTs on congruent uninformative trials (e.g., trials with high-frequency pretones and a high-frequency target) versus incongruent uninformative trials (e.g., trials with high-frequency pretones but a low-frequency target; **Table 4**). Finally, RTs were faster on uninformative trials than on informative trials (Mann-Whitney U, Monkey C, informative median (*N*=13604)=961 ms, uninformative median (*N* =14192)=646 ms, *z* =95.74, *p*<10^-06^; Monkey M, informative median (*N*=14265)=759 ms, uninformative median (*N*=20781)=459 ms, *z*=132.31, *p*<10^-06^).

In short, the monkeys’ choices and RTs depended systematically on the informative and uninformative cues. During congruent (incongruent) trials, the monkeys made more (fewer) choices toward the frequency indicated by the informative LEDs and pretones and had faster (slower) RTs.

### Informative and uninformative cues modulate evoked response to target tone burst

To understand how informative and uninformative cues modulate activity and functional connectivity in the ventral auditory pathway, which has a causal role in auditory decisions (Cohen et al., 2016; Tsunada et al., 2016, 2019), we recorded simultaneously from the AC and the vlPFC (19 sessions from monkey C and 15 from monkey M) while the monkeys performed the frequency-discrimination task (**Fig. 3**). We focused our analyses on theta-band (4–8 Hz) activity because theta oscillations have been implicated in coordinating top-down and bottom-up information flow under changing task demands (Womelsdorf et al., 2010), and causal manipulations of frontal theta indicate that this frequency band plays a key role in prioritizing and maintaining behaviorally relevant representations (Riddle et al., 2020). Consistent with these proposed functions, in both AC and vlPFC, theta-band (4–8 Hz) LFP activity was modulated by the target tone (**Fig. 4**). **Fig. 4A** shows the session-averaged evoked-response time courses of AC and vlPFC theta power, collapsed across recording sites for each monkey. In general, AC theta power was stronger than vlPFC theta power during target-tone presentation, especially on informative trials. These effects were also evident on a session-by-session basis (**Fig. 4B**, **Table 5**).

**Figure 3:**
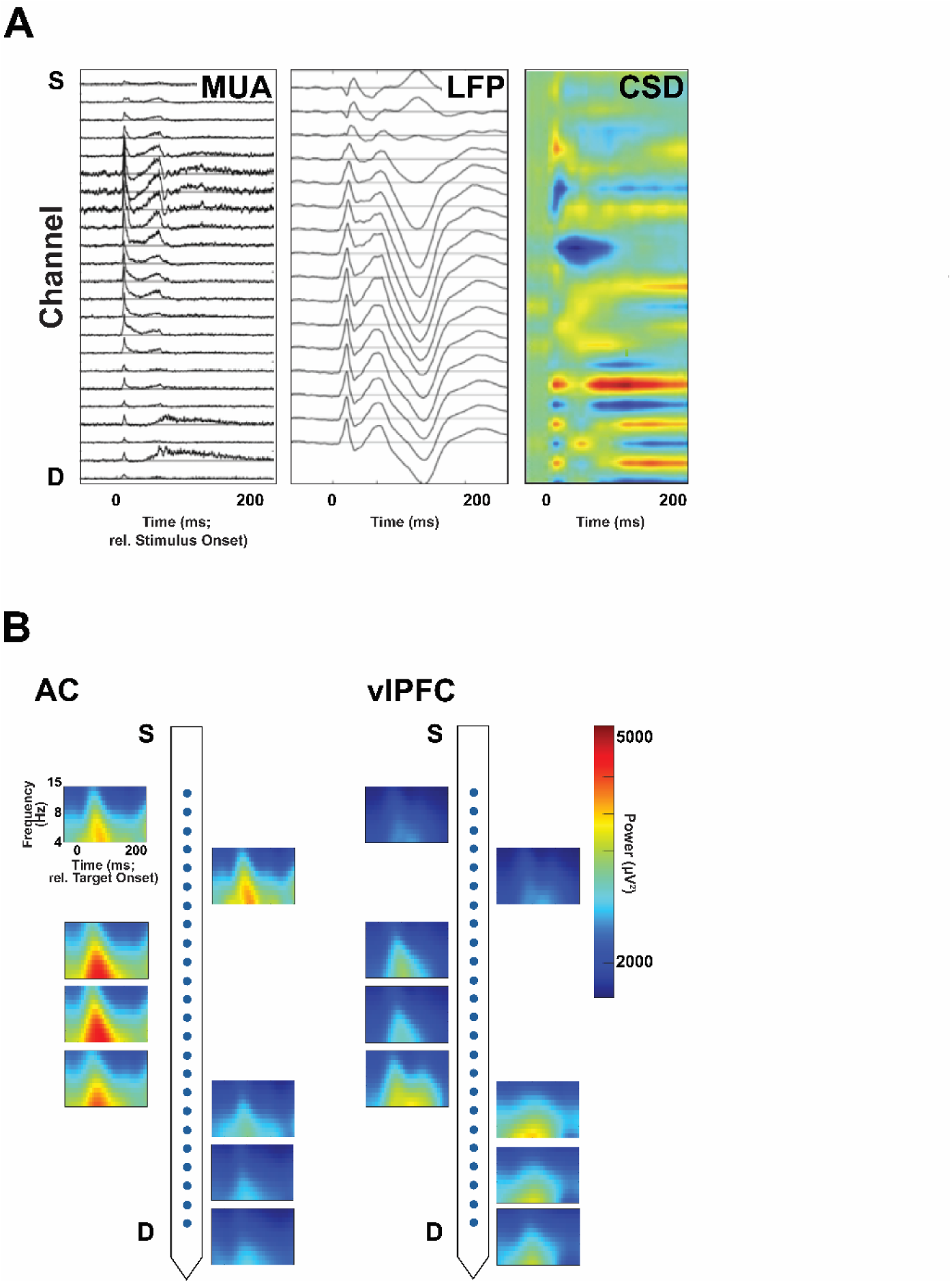
AC Laminar response profile and AC and vlPFC auditory-evoked LFPs. (**A**) Representative laminar response profile from a single recording in A1, shown as a function of channel number (channel spacing = 100 µm) from superficial (top) to deep (bottom) and time (ms) relative to onset of a broad-band auditory stimulus. **Left**, rectified multiunit activity (MUA). **Middle**, auditory-evoked potentials from the local-field potential (LFP). **Right**, two-dimensional current source density (CSD). In this color representation of the CSD, current sinks (net inward transmembrane current) are shown in blue, whereas current sources (net outward transmembrane current) are shown in red. (**B**) AC (**left**) and vlPFC (**right**) LFPs that were generated during a single recording session. In each subplot, colors indicate power at each frequency-time point (see scale at right), relative to onset of the target tone (see axis labels at upper left). LFPs are organized as a function of depth from superficial (S) recording sites (top of the electrode) to deep (D) recording sites (bottom of the electrode).

**Figure 4:**
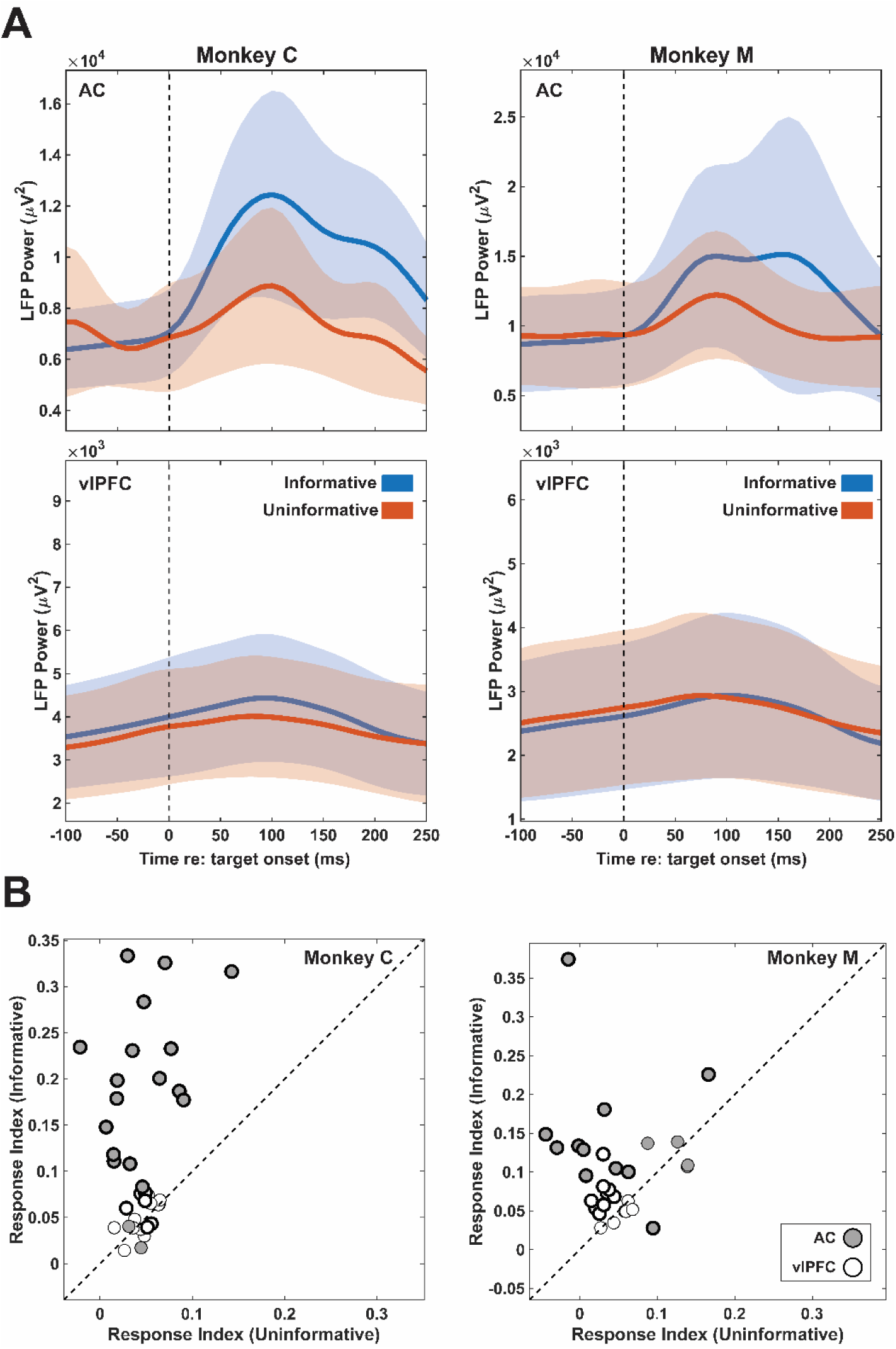
Expectation modulated AC and vlPFC LFPs. (**A**) The average time course of broadband AC (**top**) and vlPFC (**bottom**) LFP power during target-tone presentation during informative correct trials (blue data) and during uninformative correct trials (orange data). The lines and shading indicate the mean±95% confidence intervals across sessions. Note the different scales on the two y-axes. **(B)** Scatterplot of response indices on uninformative versus informative trials. Points are median index values computed for individual sessions from AC (gray) and vlPFC (white). Bold outlines denote sessions in which informative and uninformative response indices differed significantly across recording channels (paired Wilcoxon signed-rank test, p < 0.05). The dashed line is the line of equality.

### Coherence between AC and vlPFC dynamically changes over time and is modulated by expectation

To assess the impact of expectations on the functional connectivity between AC and vlPFC, we first used coherence, which quantifies the degree to which oscillatory activity in AC and vlPFC remains synchronized over a window of time. Coherence is a normalized metric that reflects the consistency of the phase relationship between the two signals rather than absolute amplitudes. Thus, two signals can exhibit high coherence even when one has a substantially larger magnitude than the other.

To provide some intuition on the relationship between coherence and LFP power, we plot in **Fig. 5A, B** the time course of AC and vlPFC LFP power during the target period from an illustrative session. During informative trials (**Fig. 5A**), the temporal dynamics of LFP power were similar in these AC and vlPFC sites. That is, soon after target onset, LFP power in both cortical areas increased, peaked around ∼75 ms, and then decreased. Reflecting these similar temporal dynamics, the coherence between these two sites was relatively high (coherence=0.15). In contrast, on uninformative trials (**Fig. 5B**), the increases/decreases in LFP power at the same two sites were not as tightly coupled, resulting in a lower coherence value (coherence=0.06). These trends were evident across our population of paired AC–vlPFC sites, where coherence tended to be stronger on informative trials than on uninformative trials (paired Wilcoxon signed-rank test, Monkey C, *N*=7600, *z*=49.76, *p*<10^-06^; Monkey M, *N*=6000, *z*=46.29, *p*<10^-06^; **Fig. 5C, D**).

**Figure 5:**
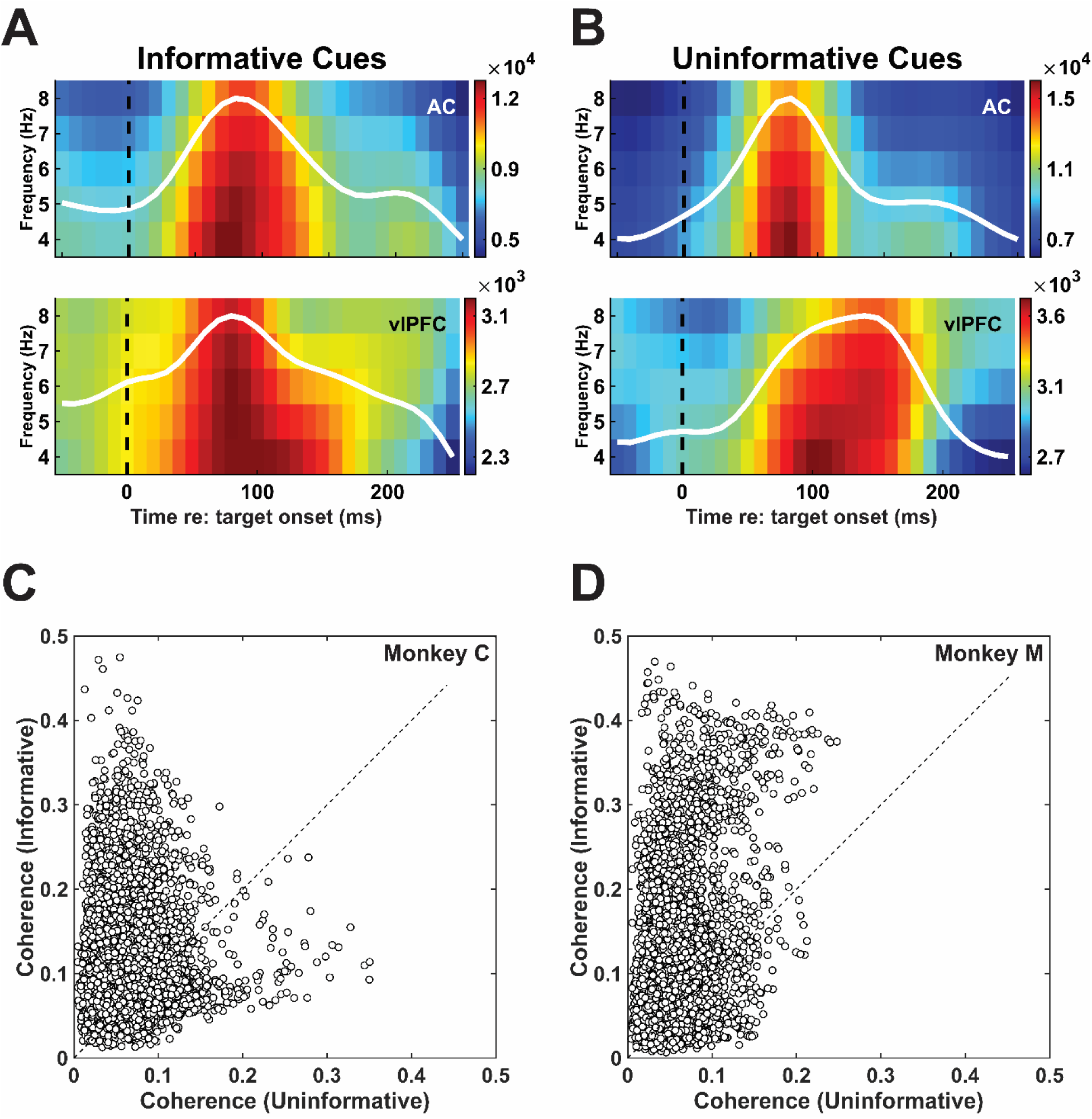
Expectation modulated AC-vlPF Coherence. (**A**) Theta-band LFP power spectrogram, relative target-tone onset during informative trials, from a single pair of AC (**top**) and vlPFC (**bottom**) recording sites. Note different color scales to the right of each panel. The average time-varying power (white line) is overlaid on each spectrogram. Data are from correct trials across all trial conditions. (**B**) Same recording sites and same format as in **A** but for uninformative trials. (**C**, **D**) Scatterplots showing the relationship between AC–vlPFC coherence during informative trials and uninformative trials for each pair of recording sites from Monkey C (**C**, N = 7600) and monkey M (**D,** N=6000) across sessions (Monkey C, 19 sessions; Monkey M, 15 sessions). Coherence magnitudes are not comparable across panels because of differences in trial subsampling.

These expectation-dependent effects on AC-vlPFC coherence changed over the course of a trial. On informative trials, coherence values were smaller during LED presentation than during the target-tone presentation (**Fig. 6**, paired Wilcoxon signed-rank test, Monkey C, *N*=7600, *z*=51.25, *p*<10^-06^; Monkey M, *N*=6000, *z*=37.14, *p*<10^-06^). In contrast, on uninformative trials, the opposite pattern emerged: coherence values were larger during the LED presentation than during target-tone presentation (paired Wilcoxon signed-rank test, Monkey C, *N* =7600, *z*=-6.09, *p*<10^-06^; Monkey M, *N*=6000, *z*=-37.90, *p*<10^-06^; **Fig. 6**).

**Figure 6:**
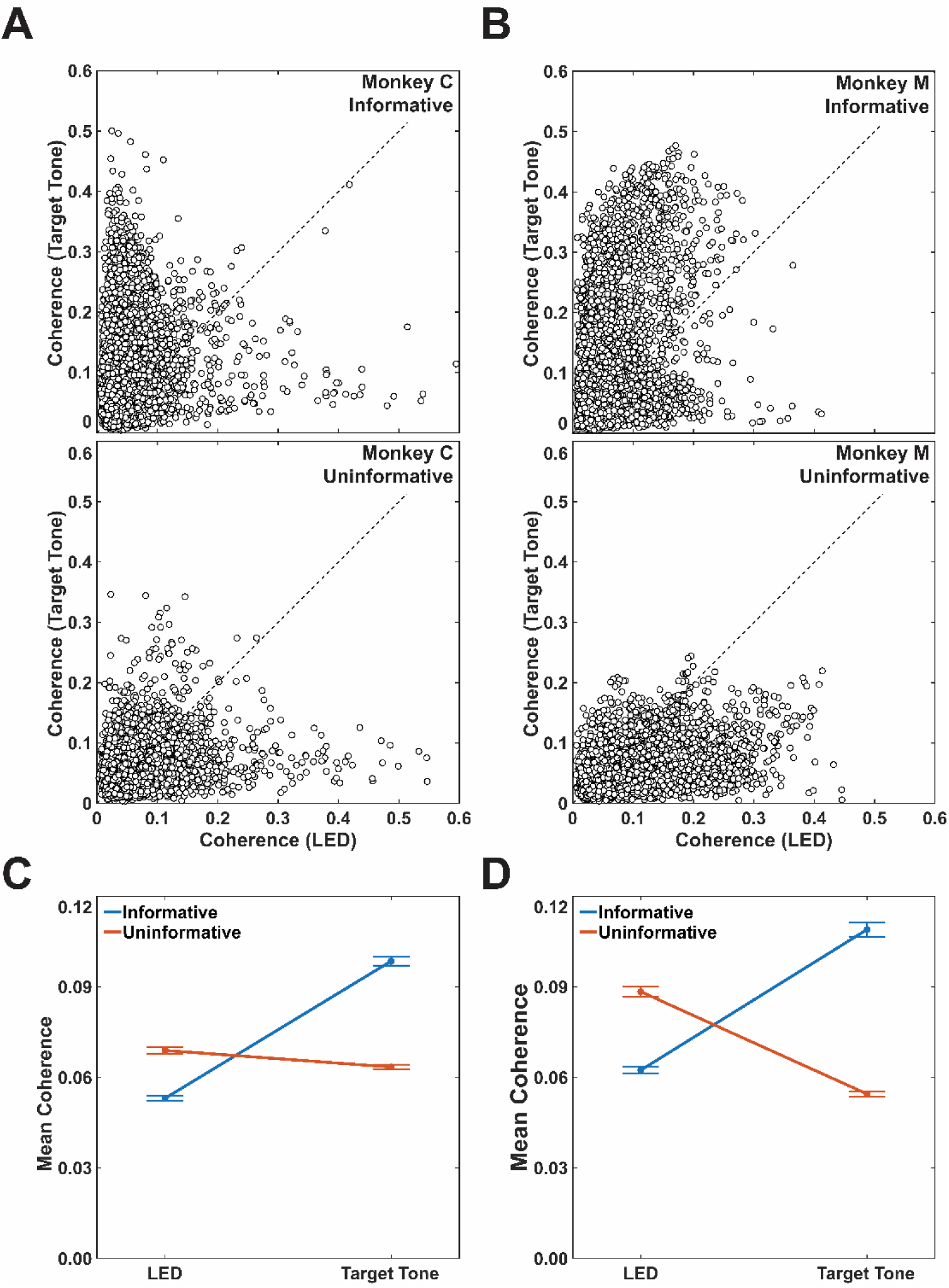
Task epoch modulated coherence. Scatterplots showing the relationship between AC–vlPFC coherence during the presentation of the target tone (x-axis) and LED (y-axis) for each pair of recording sites from Monkey C (**A**, *N*=7600 pairs, pooled across 19 sessions) and Monkey M (**B**, *N*=6000 pairs, pooled across 15 sessions). Data on the **top** row were generated during informative trials; data on the **bottom** row were generated during uninformative trials. (**C, D**) The median (points) and 95% confidence intervals (error bars) coherence values during LED and target-tone presentation for informative (blue lines) and uninformative (orange lines) trials for Monkey C (**C**) and Monkey M (**D**). Coherence magnitudes are not comparable across expectation conditions because of differences in trial subsampling; rather, note the different directions of the effects for the two conditions.

### Expectation, congruency, and SNR dynamically modulate the direction of information flow between AC and vlPFC

The coherence analyses described above measured how expectations altered AC-vlPFC functional connectivity but cannot determine the direction of this coupling. We therefore quantified the phase slope index (PSI) between AC and vlPFC sites. Unlike coherence, PSI quantifies the direction of the functional coupling by estimating the temporal lead–lag relationship between two neural signals. We defined negative and positive PSI values as “bottom-up” (AC→vlPFC) and “top-down” (vlPFC→AC), respectively. We analyzed only the site pairs that maintained a consistent PSI direction across both informative and uninformative trials (e.g., both negative or both positive for both correct and error choices; see **Fig. S1**) to focus on how task manipulations affected PSI magnitude.

We first examined how PSI values were modulated by the LED on informative versus uninformative trials. The presentation of the neutral, uninformative LED induced stronger PSI values in both directions than the informative LEDs (paired Wilcoxon signed-rank test, Monkey C, top-down, *N*=1783, *z*=12.59, *p*<10^-06^, bottom-up, *N*=1969, *z*=15.92, *p*<10^-06^; Monkey M, top-down, *N*=1965, *z*=18.18, *p*<10^-06^, bottom-up, *N*=1583, *z*=13.84, *p*<10^-06^; **Fig. 7**).

**Figure 7:**
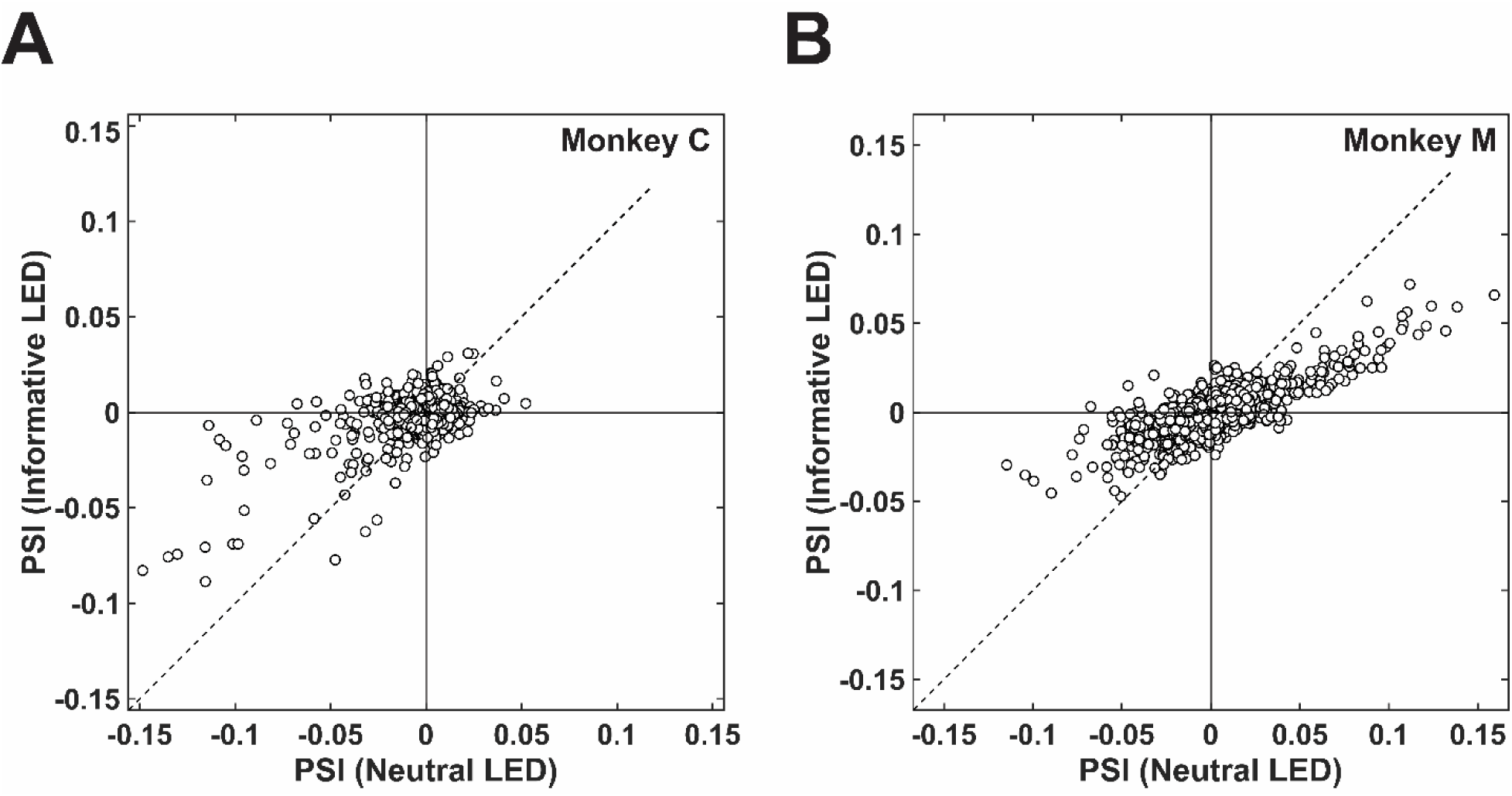
Uninformative cues induced stronger PSI values than informative cues. Scatterplots of PSI values during the presentation of the neutral LED (x-axis) and informative LED (y-axis) trials for Monkey C (**A**) and Monkey M (**B**). Each plot shows data from all the recording-site pairs from Monkey C (N=7600, pooled across 19 sessions) and monkey M (*N*=6000, pooled across 15 sessions). PSI magnitudes are not comparable across panels because of differences in trial subsampling.

Next, we examined how PSI values were modulated during target-tone presentation on informative and uninformative trials. We focused initially on low-SNR trials, when priors tend to exert the most influence on choice. On informative, congruent trials, PSI values were higher when the monkeys incorrectly chose against the prior (e.g., if the prior predicted “high frequency” and the monkey reported “low frequency”) than when their reports were correct and consistent with the prior (paired Wilcoxon signed-rank test, Monkey C, top-down, *N*=1813, *z*=16.20, *p*<10^-06^, bottom-up, *N*=2385, *z*=18.99, *p*<10^-06^; Monkey M, top-down, *N*=1761, *z*=14.11, *p*<10^-06^, bottom-up, *N*=1642, *z*=14.03, *p*<10^-06^; **Fig. 8A, B**). In contrast, on incongruent trials, PSI values were higher when the monkeys correctly chose against the prior than when they incorrectly followed the prior (Monkey C, top-down, *N*=2022, *z*=-5.28, *p*<10^-06^, bottom-up, *N*=2118, *z*=-5.44, *p*<10^-06^; Monkey M, top-down, *N*=1673, *z*=-7.99, *p*<10^-06^, bottom-up, *N*=1650, *z*=-9.32, *p*<10^-06^; **Fig. 8C, D**). These results imply stronger functional connectivity occurs when the monkeys’ expectations (generated by informative cues) are contradicted by the frequency of the target tone.

**Figure 8:**
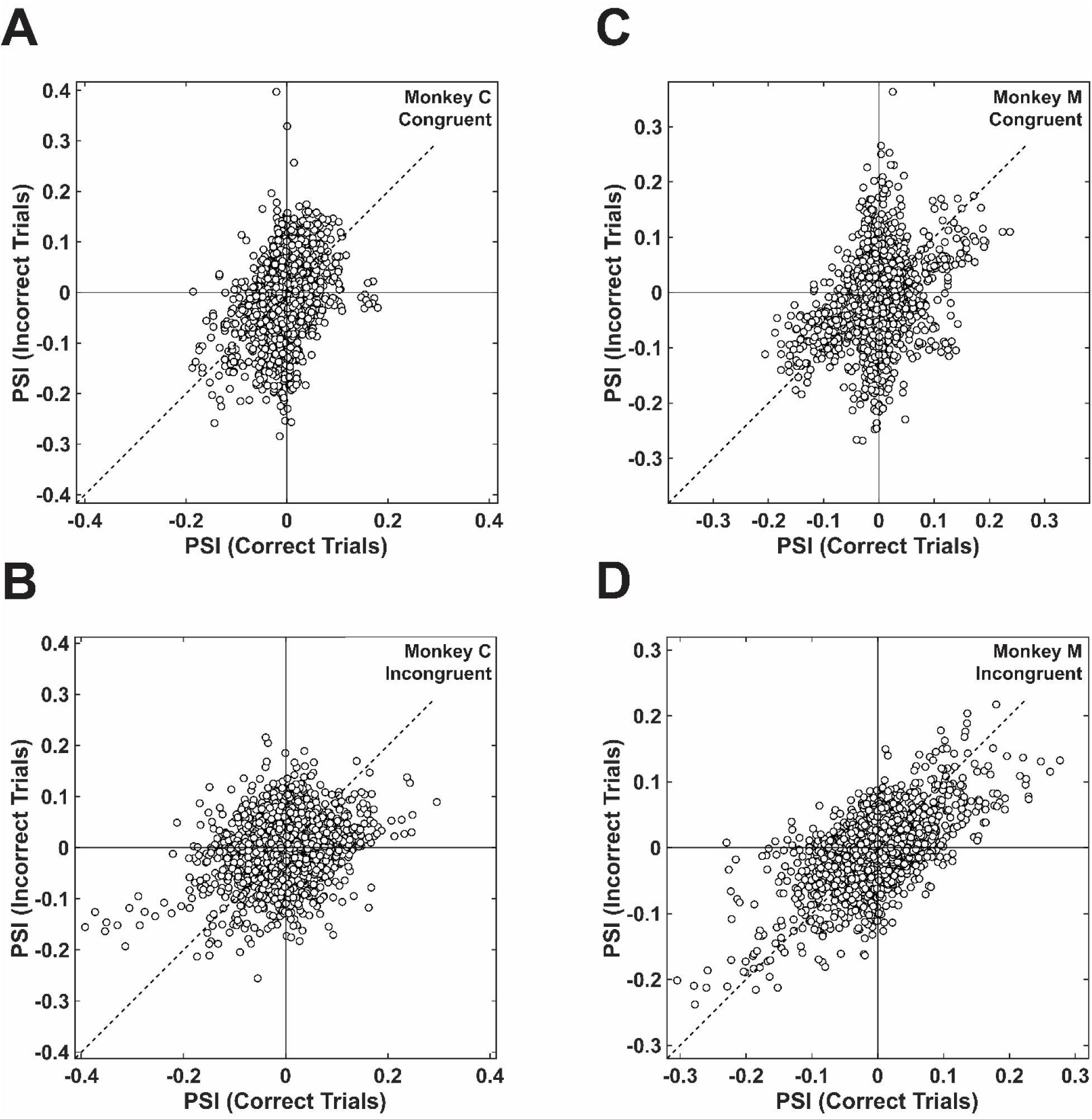
Congruency on low-SNR informative trials modulated PSI values. Scatterplots of PSI values from correct (x-axis) and error (y-axis) trials for Monkey C (**A, B**) and Monkey M (**C, D**) as a function of congruent (the prior aligns with the frequency of the target tone; **top** row) and incongruent (the prior conflicts with the frequency of the target tone; **bottom** row) trials. Each plot shows data from all the recording-site pairs from monkey C (N=7600, pooled across 19 sessions) and monkey M (*N*=6000, pooled across 15 sessions). PSI magnitudes are not comparable across panels because of differences in trial subsampling.

On uninformative low-SNR trials, PSI was less sensitive mismatches between expectations based on the pretones and the target tone. On congruent trials, PSI values were larger when the monkeys made errors and went against the prior but only in the top-down direction (Monkey C, top-down, *N*=1982, *z*=10.14, *p*<10^-06^; Monkey M, top-down, *N*=1376, *z*=8.52, *p*<10^-06^). In the bottom-up direction, the relationship was statistically reliable only for Monkey C (Monkey C, bottom-up, *N*=1912, *z*=5.32, *p*<10^-06^; Monkey M, bottom-up, *N*=1657, *z*=1.72, *p*=0.09; **Fig. 9A, B**). On incongruent trials, we could not identify any effect of choice on PSI (Monkey C, top-down, *N*=1858, *z*=-1.21, *p*=0.228, bottom-up, *N*=2026, *z*=2.15, *p*=0.03; Monkey M, top-down, *N*=1502 *z*=1.34, *p*=0.18, bottom-up, *N*=1470, *z*=1.20, *p*=0.23; **Fig. 9C, D**).

**Figure 9:**
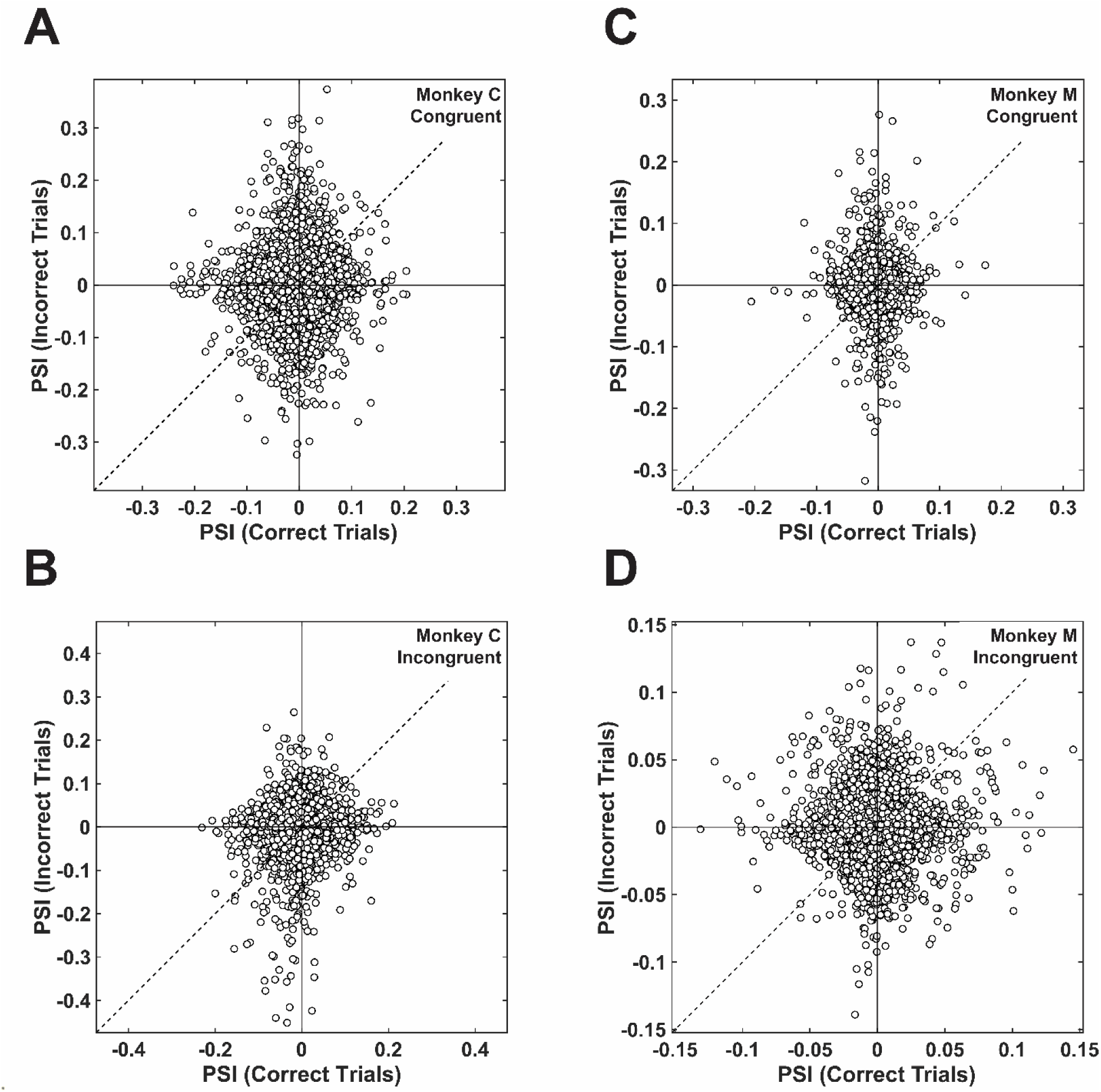
Congruency on low-SNR uninformative trials modulated PSI values. Scatterplots of PSI values from correct (x-axis) and error (y-axis) trials for Monkey C (**A, B**) and Monkey M (**C, D**) as a function of congruent (the prior aligns with the frequency of the target tone; **top** row) and incongruent (the prior conflicts with the frequency of the target tone; **bottom** row) trials. Each plot shows data from all the recording-site pairs from monkey C (N = 7600, pooled across 19 sessions) and monkey M (*N* = 6000, polled across 15 sessions). PSI magnitudes are not comparable across panels because of differences in trial subsampling.

A different pattern emerged on high-SNR trials. Error trials tended to generate stronger PSI values than correct trials on both informative (**Fig. S2**, **Table 6**) and uninformative (**Fig. S3**, **Table 6**) trials. Thus, AC-vlPFC connectivity was dynamically modulated by task demands, including the cue type, its relationship to the sensory information (target frequency and SNR), and whether the choice was correct or not.

### Spatial distribution of bottom-up and top-down interactions are modulated by expectation

Taking advantage of the fact that we placed our electrodes orthogonal to the cortical lamina, we assessed lamina-dependent spatial organization of PSI values. Because there is a known laminar organization of ascending and descending connectivity (Hackett et al., 1999; Romanski et al., 1999a; Romanski et al., 1999b), we tested whether putative bottom-up (AC→vlPFC) and top-down (vlPFC→AC) interactions are spatially clustered according to these innervation patterns. In particular, we organized PSI values within a 20⨉20 grid of AC–vlPFC recording sites as a function of recording depth (because each electrode had 20 channels). For each grid, we calculated Moran’s I (Moran, 1950), which is a bounded measure of spatial autocorrelation (−1 to 1, where −1 is perfect dispersion, 0 is random patterning, and 1 is perfect clustering). We quantified the spatial organization between expectation conditions by calculating the Pearson correlation coefficient as a function of grid location and then compared this observed correlation value to a null distribution generated by randomly permuting PSI values across spatial locations.

During LED presentation, two spatial patterns emerged, regardless of trial type (**Fig. 10**). First, vlPFC channels had broad top-down functional connectivity with the deep AC channels; weakly in Monkey C and more strongly in Monkey M. Second, middle AC channels showed bottom-up functional connectivity with several middle vlPFC channels. The spatial clustering of PSI values was the same on informative and uninformative trials (paired Wilcoxon signed-rank test, H_0_: session-wise Moran’s I distribution are equal; Monkey C, *N* =19, *z*=-0.12, *p*=0.90; Monkey M, *N*=15, *z*=-1.82, *p*=0.07; **Fig. 10B, C**).

**Figure 10:**
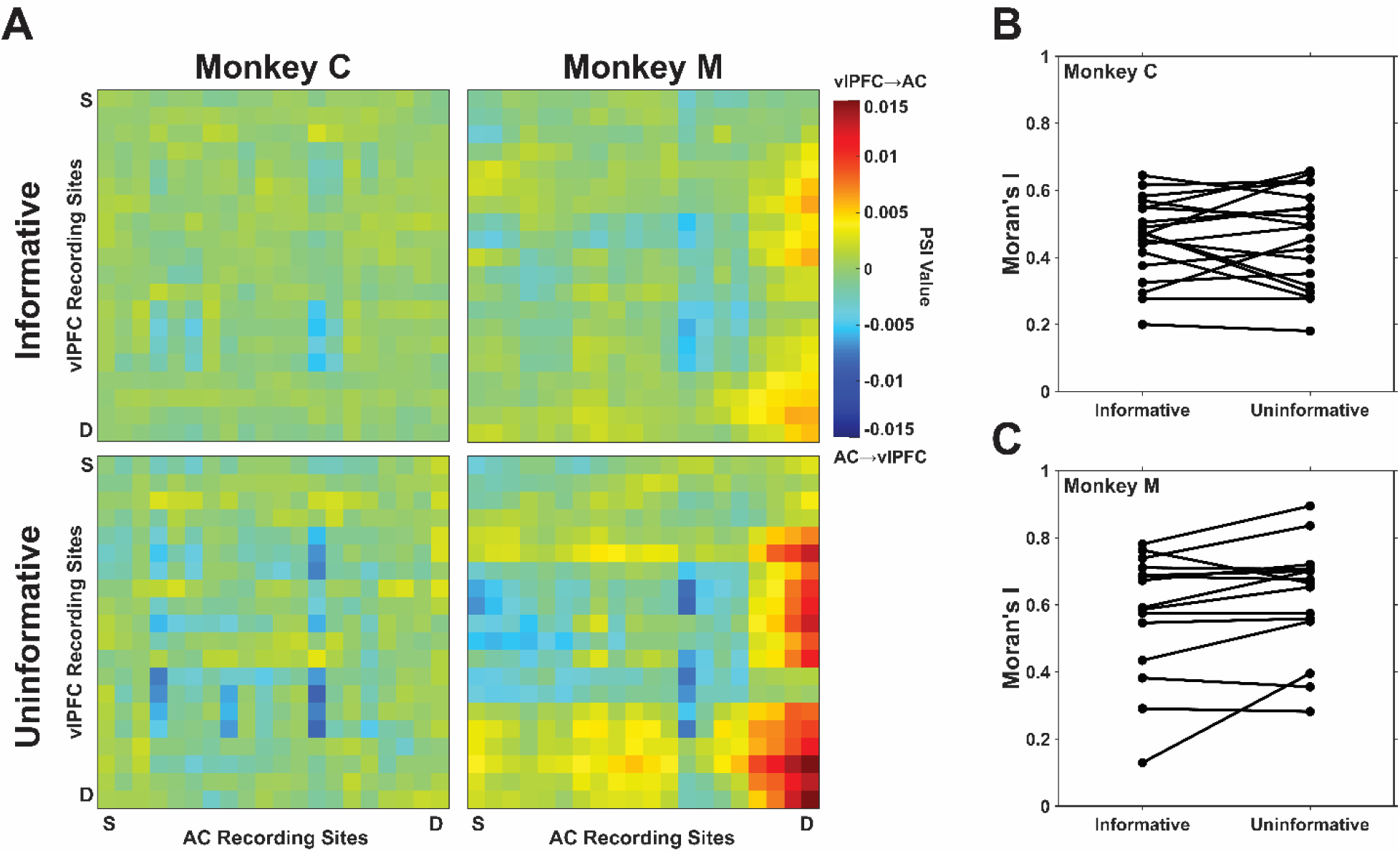
During LED presentation, expectations affected PSI magnitude but not the spatial structure of AC-vlPFC interactions. Across-session mean PSI values (Monkey C, *N*=19, Monkey M, *N*=15) as a function of each recording-site pair. These pairs are organized relative to each AC and vlPFC probe’s position in the cortex, from superficial (S) to deep (D). Positive PSI values are shown as warm colors and indicate vlPFC→AC interactions, whereas negative PSI values are shown as cool colors and indicate AC→vlPFC interactions. The spatial organization of PSI values during informative trials are plotted in the **top** row; the spatial organization of PSI values during uninformative trials are plotted in the **bottom** row. (**B, C**). Session-by-session Moran’s I values for informative and uninformative trials. Moran’s I values from the same session are connected by a line. A Moran’s I value of −1 indicates that the spatial organization of the data is dispersed, 0 indicates randomness, and 1 indicates clustered.

During target-tone presentation, the spatial pattern of AC-vlPFC connectivity was modulated by expectation and choice outcomes (**Fig. 11**). Overall, we observed a columnar pattern, with superficial and deep AC channels having broad functional connectivity with all of the vlPFC channels. In addition, vlPFC had broad functional connectivity with the middle AC channels. On correct trials, the informative trials generated stronger spatially clustering than uninformative trials (paired Wilcoxon signed-rank test, H_0_:session-wise Moran’s I distribution are equal; Monkey C, *N* =19, *z*=3.46, *p*=5.3⨉10^-4^; Monkey M, *N*=15, *z*=2.10, *p*=0.04; **Fig. 11B, C**). Correct trials also corresponded to stronger clustering than error trials on informative, high-SNR, congruent trials (Monkey C, *N*=19, *z*=3.70, *p*=2.13 ⨉ 10^-4^; Monkey M, *N*=15, *z*=2.61, *p*=8.9⨉10^-3^; **Fig. S4**).

**Figure 11:**
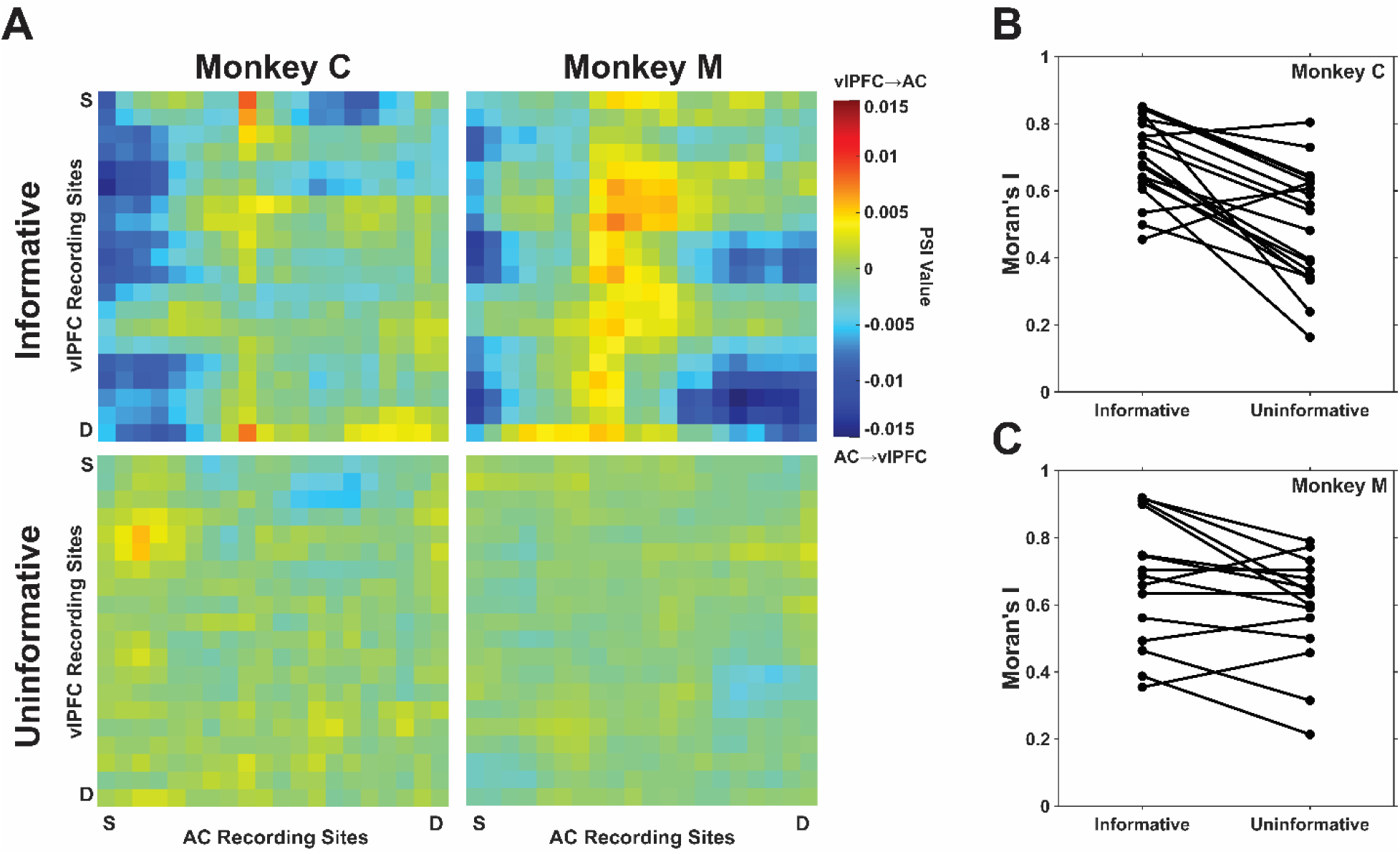
During target-tone presentation, expectations affected PSI spatial organization. (**A**) Across-session mean PSI values (Monkey C, *N*=19, Monkey M, *N*=15) during the target-tone period as a function of each recording-site pair. These pairs are organized relative to each AC and vlPFC probe’s position in the cortex, from superficial (S) to deep (D). Positive PSI values are shown as warm colors and indicate vlPFC→AC interactions, whereas negative PSI values are shown as cool colors and indicate AC→vlPFC interactions. The spatial organization of PSI values during informative trials are plotted in the **top** row; the spatial organization of PSI values during uninformative trials are plotted in the **bottom** row. (**B, C**). Session-by-session Moran’s I values for informative and uninformative trials. Moran’s I values from the same session are connected by a line. A Moran’s I value of −1 indicates that the spatial organization of the data is dispersed, 0 indicates randomness, and 1 indicates clustered.

## Discussion

We tested how different sources of expectations shaped behavior and bidirectional AC-vlPFC functional connectivity during auditory decision-making. Informative rule-based cues biased choices in the direction of the cue, with faster responses when the cue and stimulus were congruent but slower responses when they were incongruent. Uninformative stimulus-based cues had similar, albeit smaller and more variable, behavioral effects. Despite these similar behavioral effects, the two cue types differentially modulated AC–vlPFC connectivity both before the decision, when expectations were being established, and while the decision was being formed.

### LED Cues Differentially Modulate AC-vlPFC Functional Connectivity

Before the decision was formed, AC-vlPFC connectivity tended to be stronger on trials with uninformative versus informative cues. These differences did not correspond to changes in the spatial organization of top-down and bottom-up interactions within the AC–vlPFc circuit (**Figs. 7, 10**). Together, these results suggest that the same AC and vlPFC sites remained engaged across expectation conditions but their functional connectivity was modulated by the source of the expectation.

A priori, we had expected the opposite result: because the informative LEDs cued specific rules associated with the eventual target, these LEDs would induce more AC-vlPFC communication than the uninformative LEDs. One possible explanation for our findings is that, following the informative LEDs, the upcoming sensory input is more expected, so less information needs to be exchanged between AC and vlPFC prior to the target tone. This explanation is similar to the idea that expertise or learning often reduces large-scale recruitment as local processing becomes more efficient (Alain et al., 2007; Jensen & Mazaheri, 2010; Weisz et al., 2014). Alternatively, the neutral LED might have induced an anticipatory or preparatory state (Brosch et al., 2005; Fritz et al., 2007; Guo et al., 2019; Mohn et al., 2021; Yin et al., 2008) that resulted in stronger functional connectivity. Consistent with this idea of increased preparation, RTs tended to be faster on uninformative than informative trials.

### AC-vlPFC Connectivity During Decision Formation Depends on Expectations and Task Demands

Although increased connectivity has been interpreted as predictive of improved behavioral performance (Fries, 2015; Siegel et al., 2012), our findings suggest that stronger AC-vlPFC does not simply reflect better behavior. Instead, we found that connectivity strength is more closely aligned with expectation-sensory conflict than with behavioral performance itself (e.g., **Figs. 8, 9**), which was especially true on low-SNR trials that should benefit most from prior information. Further, the effects of expectations on these connectivity patterns modulated not only their strength but also their spatial organization.

This pattern of similar behavioral reports with different underlying neural correlates is consistent with our previous findings that used a similar task design with human participants (Tardiff et al., 2022). In that work, informative, rule-based expectations and uninformative, stimulus-based expectations had superficially similar effects on choice and RT behavior but distinguishable computational (effects on fit parameters from a drift-diffusion model) and physiological (pupil responses that were sensitive to violations of expectations for the informative but not uninformative, cues) signatures. More work is needed to tie those signatures more directly to the changes in functional connectivity that we identified in our current study.

### Where are the directional effects?

Despite strong anatomical predictions about the direction of information flow within the AC– vlPFC circuit (Chennu et al., 2013; Hannemann et al., 2007; Park et al., 2015; Patel et al., 2022; Zekveld et al., 2006), we observed relatively few robust directional effects during either of the two epochs. For example, we reasoned that when the monkeys were presented informative cues, there should be a top-down bias corresponding to strong top-down connectivity between superficial/deep vlPFC neurons and deep/superficial AC neurons (Kaas & Hackett, 1998; Romanski et al., 1999a; Romanski, et al., 1999b). We did observe columnar, top-down signaling from vlPFC arriving in the deep layers of the AC (**Fig. 10**), consistent with anatomical innervation. However, both positive and negative PSI values tended to be weaker on informative versus uninformative trials (**Fig. 7**). Similarly, we did not observe any consistent directional bias during decision formation at the population level in either expectation condition (**Fig. 8, 9**).

Qualitatively, the spatial distribution of bottom-up interactions on informative trials was broadly consistent with known corticocortical anatomy, with superficial/deep AC neurons projecting to deep/superficial vlPFC neurons. In contrast, the spatial organization of top-down interactions was less consistent with anatomical predictions, with vlPFC→AC communication appearing to terminate within the middle layers of A1 rather than the deep layers where cortico-cortico feedback projections are typically observed (Van Essen & Fellerman, 1991). Instead, this pattern is more reminiscent of ascending innervation patterns of neuromodulatory systems like the cholinergic (Disney et al., 2012) system that are known to alter neural responses to ascending thalamic signals. It seems plausible that feedback mechanisms that serve auditory processing may also target the thalamo-cortico recipient layer in this way.

However, interpreting these spatial maps as functional overlays to known anatomical connections should be done with caution, for several reasons. First, because our recording sites are not connected monosynaptically, it may not be possible to map schematized patterns of neuroanatomy to our functional connectivity patterns. Second, because we were analyzing relatively slow (theta) oscillations, we may be observing longer steady-state activity that involves many top-down and bottom-up loops of neural activity. Third, rule-based and stimulus-based expectations may not map cleanly onto a simple top-down versus bottom-up cortical neuroanatomy. To resolve these issues, we would need to measure more spatially and temporally precise neural signals, such as spike-field coherence in which we can conceptualize spikes as the output of a neural process whereas the LFPs are the input to neurons (Logothetis, 2003; Logothetis et al., 2001; Mitzdorf, 1985). Spike–field coupling would therefore provide a more direct, anatomically anchored test of our LFP findings and link the observed theta-band connectivity with pathway-specific changes in local spiking output.

## Conclusion

This study demonstrates that similar expectation-dependent behavioral biases can emerge from distinct functional communication states within the AC–vlPFC circuit. Expectation influenced not only the magnitude of functional connectivity, but also its spatial organization and dependence on task demands. Whereas previous studies of the ventral auditory pathway have focused primarily on single-unit activity, our findings reveal how populations of neurons coordinate across AC and vlPFC. By characterizing both coherence and PSI between LFP signals, this work provides a circuit-level framework for understanding how different sources of expectation dynamically modulate auditory processing during decision-making.

## Materials and Methods

### Subjects and Surgical Procedures

All procedures and experiments conformed to the National Institutes of Health Guide for the Care and Use of Laboratory Animals and were approved by the University of Pennsylvania Institutional Animal Care and Use Committee. Two adult male monkeys (*Macaca Mulatta*, Monkey C, 8 years old, and Monkey M, 12 years old) participated in this study. All surgical procedures were conducted under general anesthesia and used aseptic surgical techniques. Before each surgery, we identified the stereotactic locations of the auditory cortex (AC) and ventrolateral prefrontal cortex (vlPFC) through anatomical MRI scans (Christison-Lagay et al., 2017; Cohen et al., 2004; Johnston et al., 2016; Romanski & Goldman-Rakic, 2002). Using these scans, we designed a titanium recording chamber that: (1) allowed simultaneous access to both cortical fields, and (2) was oriented perpendicular to the cortical layers (Frey et al., 2011). Further details on the surgical approach can be found in (Johnston et al., 2016). We placed the recording chambers over the right hemisphere in Monkey C and over the left hemisphere in Monkey M.

### Auditory Frequency-Discrimination Task

In each session, the monkey was seated in a primate chair that was located in an RF-shielded room; this room had sound-attenuating walls and echo-absorbing foam on the inner walls. A calibrated free-field speaker (MSP7, Yamaha) was placed 1 m in front of the monkey and at their eye level. A cluster of green, blue, and yellow equiluminescent LEDs was centered on the speaker face. The monkey moved a joystick, which was attached to the primate chair, to indicate whether a target tone burst (200-ms duration; 65 dB SPL; 5-ms cos^2^ ramp) was “low frequency” or “high frequency” (**Fig. 1**). We titrated task difficulty by embedding the target tone in a broadband background noise (0.1-16 kHz). We varied the sound level of the target tone between 60 and 85 dB SPL. As a result, the signal-to-noise ratio (SNR) was between −5 and +20 dB. On some trials, we presented the background noise alone; on these trials, the monkey was randomly rewarded, independent of their choice.

Because we were interested in how expectations modified the monkey’s decisions, we presented either rule-based (“informative”) cues or stimulus-based (“uninformative”) cues before target-tone presentation. Informative trials started with the onset of a colored LED (200-ms duration). The LED color served as an informative prior: a blue or green LED indicated a 75% chance of a high-frequency or low-frequency target tone, respectively. Following the offset of the LED and a silent period (600 ms), the target tone burst was presented. The color of the LED was constant within a block of trials but changed randomly across trial blocks.

Uninformative trials began with a yellow LED, which provided no predictive information. After a random interval of 250–600 ms, we presented three tone bursts (e.g., “pretones”). The frequencies of all three pretones (200-ms duration; 65 dB SPL; 5-ms cos^2^ ramp) were the same (i.e., either all “low frequency” or all “high frequency”).

The frequencies of the tone bursts were based on the best frequency of the auditory-cortex recording sites (see below). Each session followed an alternating block design, with the best frequency used as either the low or frequency in the given block. When the best frequency was the low (high) frequency, the high (low) frequency was approximately 1-log step above (below).

### Statistical Analysis of Expectation on Choice and Response Time

We quantified the effect of the informative LEDs on choice behavior by fitting a binomial generalized linear mixed-effects model with a logit link. Choice was coded as a binary response (“high” choice=1; “low” choice=0), informative LEDs were treated as a categorical fixed effect (with the neutral LED as the reference level), and SNR was a continuous covariate. All models included a random intercept for session to account for repeated measurements within sessions. We fit a single GLMM spanning all trials with the fixed-effects structure Choice ∼ LED × SNR + (1|Session).

To examine specifically the effect of the (uninformative) pretones on choice, we generated a modified GLM to evaluate choice. This model included pretone identity (“high-frequency pretones”=+0.5, “low-frequency pretones”=−0.5), choice (“high-frequency” choice=1; “low-frequency” choice=0), and SNR as a continuous covariate. The model took the following form: Choice ∼ Pretone × SNR + (1|Session).

We limited our analyses to RTs generated only from correct trials and excluded SNR=0 trials because there was not a correct answer on these trials. We categorized trials as a function of whether the LED was neutral, “congruent” with the target (e.g., blue LED cue with a high-frequency target), or “incongruent” with the target (e.g., blue LED with a low-frequency target). Our model had the form of: RT ∼ Congruency + |SNR| + (1|Session). Similarly, for testing the effect of pretones on RTs, we categorized trials by pretone–target congruency (e.g., a congruent trial had high-frequency pretones with high-frequency target; incongruent trials had high-frequency pretones with a low-frequency target). The model took the form of: RT ∼ Congruency + |SNR| + (1|Session).

### Recording Strategy

During each recording session, we placed one 24-channel electrode (Plexon S-Probes, Plexon, Inc., Dallas, TX, USA; an inter-channel spacing of 100 µm) in the AC and a second one in the vlPFC. Neural signals were amplified (PZ5, Tucker-Davis Technologies), digitized, and stored (sampling rate: 24.4 kHz; RZ2, Tucker-Davis Technologies) for online and offline analyses.

While the AC electrode was advanced, we presented an auditory search stimulus (80-ms white-noise burst at 65 dB SPL) to identify evoked responses. Once we found auditory responses, we generated the site’s current-source-density profile from ∼150 repetitions of the search stimulus. For the auditory-cortex recordings, we adjusted the electrode depth to ensure that the highest rate of multiunit activity and the initial short-latency current sink were aligned with the middle channels of the electrode. This sink represents current flow associated with the neural circuitry of the thalamorecipient zone (Freeman & Nicholson, 1975; Mackey et al., 2024; Nicholson & Freeman, 1975; Reser et al., 2000). Next, we retracted the electrode by 250–500 μm and allowed the tissue to stabilize for >50 mins to minimize electrode drift. Because the laminar profile of neural activity in the vlPFC is not known, we opted for a different approach and positioned the vlPFC electrode so that its most superficial recording site had auditory-evoked multi-unit activity. This ensured the probe’s contacts spanned all cortical layers.

Next, monkeys listened passively to a dynamic moving ripple (DMR) noise. We generated a site’s spectrotemporal response field (STRF) from the single-unit activity collected during this period. While continuing to record neural activity, the monkey participated in blocks of trials of the frequency-discrimination task. After this behavioral-recording session was concluded, we once again presented the search stimulus and DMR stimuli and regenerated the current-source-density profile and the STRFs to ensure that the recording site was stable. We included only stable recordings in this study.

### Dynamic moving ripple (DMR) noise: stimulus to generate a neuron’s STRF

The DMR noise (Escabí & Schreiner, 2002; Miller et al., 1996) was a continuous time-varying broadband noise stimulus that spans 0.1–32 kHz (10-min duration; 65 dB spectrum level per ⅓ octave; 96 kHz sampling rate; 24-bit resolution). At any instant of time, the stimulus had a sinusoidal spectrum; the density of the spectral peaks was determined by the spectral modulation frequency ( = 0–4 cycles/octave). The peak-to-peak amplitude of the ripple was 30 dB. The stimulus also contained temporal modulations that were controlled by the temporal modulation frequency (0–25 Hz). Both the spectral and temporal parameters were varied randomly and dynamically; the maximum rates of change were 0.25 Hz and 1 Hz, respectively. This variability in the DMR’s spectrotemporal properties provided an opportunity to probe the spectrotemporal acoustic space in a minimally biased manner.

For each neuron, we derived its STRF using the reverse-correlation method, which is the average spectrotemporal stimulus envelope immediately preceding each spike. We used a two-step procedure to identify each STRF’s reliable samples and its overall reliability. To identify the statistically reliable spectrotemporal samples in each STRF, we generated a null model under the assumption that spiking activity occurs randomly with respect to the DMR. We calculated this null model by first randomizing the inter-spike intervals and then averaging the spectrotemporal envelope of the DMR relative to the time of each spike. This procedure generated a "noise" STRF. We repeated this procedure 1000 times to generate a distribution of spectrotemporal noise-STRF samples. A STRF sample was considered significant (*p*<0.001) if its value was outside of the 99.9% confidence interval of its respective noise-STRF sample. STRF samples that overlapped with the noise-STRF distribution were assigned a value of 0.

For any given STRF, ∼230 STRF samples (i.e., 288 temporal × 800 spectral samples × 0.001) are false positives. Although these samples exceeded our significance criterion, they did not fulfill our primary objective of identifying STRFs with reproducible and reliable auditory responses. To identify such STRFs, we computed a reliability index to identify a STRF with reproducible structure. We calculated this index by breaking the DMR stimulus into twenty 30 s long segments. We next randomly selected two subsamples of the 20 segments, generated a STRF from each of these segments, and calculated their Pearson correlation coefficient. A correlation coefficient ∼1 indicates that a STRF was very reliable, consistent with a stable stimulus-evoked and synchronized response, whereas a value ∼0 indicated that it was not reliable. We repeated this procedure 500 times to generate a distribution of correlation-coefficient values. Finally, we generated a null distribution of coefficient values using the same process but with randomized inter-spike intervals. A Mann-Whitney test evaluated the hypothesis that the actual and the null distributions of correlation coefficients had the same median values. If the null hypothesis was rejected (*p*<0.01), we considered a STRF to be significant. A STRF’s “best frequency” (BF) was the frequency value that consistently elicited the largest neural response.

### Preprocessing of Neural Data

We conducted spectral analyses on the FieldTrip (Oostenveld et al., 2011) and MATLAB (The Mathworks) platforms. After aligning neural data relative to a task event (e.g., LED onset), we downsampled (1000 Hz) and then referenced the data. We constructed referenced channels because it generates a more localized, higher-resolution neural signal by removing local sources shared across multiple channels (Bastos et al., 2020). We referenced channels by subtracting any given channel’s data from data collected at the channel 400 μm beneath it (e.g., referenced channel #1 was calculated by subtracting the actual channel #5 from the actual channel #1). This produced 20 referenced channels. We estimated the LFP power of the 200-ms task epochs (which were zero padded to 1 s) via a single-taper FFT and a smoothing window of 4 Hz.

### Quantification of Target-Tone Evoked Potentials

On a session-by-session basis, we calculated a response index: (*stim−pre*)*/*(*stim+pre*), where *stim* is theta-band power during the 250-ms period after target onset and *pre* is theta-band power during the 100-ms period preceding target onset). As a response-index value approaches +1 (−1), it implies more theta-band power during (preceding) the target tone.

### Analysis of Functional Connectivity: Coherence and Phase Slope Index

To test whether the AC-vlPFC functional connectivity was modulated by our task variables, we calculated the coherence (*Coh*), which is a non-directional measure of phase consistency, between two neural time series: 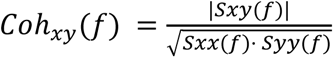, where *Sxx* and *Syy* are the auto-spectral power densities and *Sxy* is the cross-spectral power density.

A coherence value of 0 implies that there is not a linear relationship between the two time series, whereas a value of 1 implies a perfect linear relationship. Coherence reflects the consistency of the phase and amplitude relationship between two signals and is independent of their absolute amplitudes. Thus, two signals can exhibit high coherence even if their amplitudes differ substantially, provided that their phase relationship remains stable. Because coherence magnitude is dependent on trial number, we subsampled trials across experimental conditions to generate directly comparable coherence values. We subsampled the data 1000 times and reported the average value.

Phase-slope index (PSI), unlike coherence, quantifies the direction of information flow between AC-vlPFC channel pairs. PSI is based on the consistency of the phase delay between the neural signals measured at two sites (*x* and *y*): 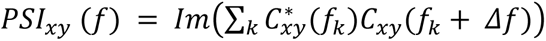, where *Im* denotes the imaginary part of the complex conjugation of coherence 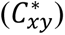, *f_k_* indexes the frequency range, and *Δf* is the bin step. A PSI value >0 indicates that *x* leads *y*. Conversely, a PSI value <0 indicates that *y* leads *x*. We calculated PSI using a *Δf* of 2 Hz. This choice preserved frequency specificity within the narrow 4-8 Hz band, at the cost of relying on fewer neighboring frequency bins and therefore produced a more conservative estimate of directional coupling. The magnitude of the absolute value of PSI reflects the strength/consistency of that directional trend. Because the magnitude of PSI, like coherence, depends on the number of trials used to compute it, we again subsampled trials across experimental conditions to generate comparable PSI values.

## Supporting information

2026_07_20_Area_Communication_Paper_Supplemental_Materials

## Data Availability

The data analyses were performed in Matlab and are currently available on GitHub (https://github.com/CohenAuditoryLab/Auditory_Functional_Communication_Toolbox) and will be updated following review and revision. Data necessary to generate the figures of this manuscript have been deposited into a stable repository (https://datadryad.org/) and will be retained for at least a 10-year period following final revision. Further inquiries concerning data and analysis code should be directed to the corresponding author.

## Additional Information

### Author Details

**Corey Roach**

**Contribution:** Conceptualization, Data Analysis, Investigation, Visualization, Methodology, Writing-original draft, Writing-review and editing

**Competing Interests:** No competing interests declared

**Lalitta Suriya-Arunroj**

**Contribution:** Conceptualization, Data Acquisition, Methodology

**Competing Interests:** No competing interests declared

**Sonia Bilal**

**Contribution:** Conceptualization, Investigation, Data Analysis

**Competing Interests:** No competing interests declared

**Sophia Fu**

**Contribution:** Conceptualization, Investigation, Data Analysis

**Competing Interests:** No competing interests declared

**Yale Cohen**

**Contribution** Conceptualization, Software, Formal analysis, Funding acquisition, Visualization, Methodology, Writing—review and editing

**Competing interests** No competing interests declared

**Joshua I Gold**

**Contribution** Conceptualization, Software, Formal analysis, Funding acquisition, Visualization, Methodology, Writing—review and editing

**Competing interests** Senior Editor, *eLife*

## Funding

This work was supported by the Army Research Laboratory (ARL-BAA-0016)

## Acknowledgments

We thank Bijan Pesaran and Agrita Dubey for their vital feedback during the early conceptualizations of our main analyses. We’d also like to thank Harry Shirley and ULAR for providing the outstanding primate husbandry throughout the study.

## Notes

### Competing Interest Statement

The authors have declared no competing interest.

## References

Alain, C., Snyder, J. S., He, Y., & Reinke, K. S. (2007). Changes in auditory cortex parallel rapid perceptual learning. Cerebral Cortex (New York, N.Y.: 1991), 17(5), 1074–1084. 10.1093/cercor/bhl018

Bastos, A. M., Lundqvist, M., Waite, A. S., Kopell, N., & Miller, E. K. (2020). Layer and rhythm specificity for predictive routing. Proceedings of the National Academy of Sciences of the United States of America, 117(49), 31459–31469. 10.1073/pnas.2014868117

Brosch, M., Selezneva, E., & Scheich, H. (2005). Nonauditory events of a behavioral procedure activate auditory cortex of highly trained monkeys. The Journal of Neuroscience: The Official Journal of the Society for Neuroscience, 25(29), 6797–6806. 10.1523/JNEUROSCI.1571-05.2005

Chennu, S., Noreika, V., Gueorguiev, D., Blenkmann, A., Kochen, S., Ibáñez, A., Owen, A. M., & Bekinschtein, T. A. (2013). Expectation and attention in hierarchical auditory prediction. The Journal of Neuroscience: The Official Journal of the Society for Neuroscience, 33(27), 11194–11205. 10.1523/JNEUROSCI.0114-13.2013

Christison-Lagay, K. L., Bennur, S., & Cohen, Y. E. (2017). Contribution of spiking activity in the primary auditory cortex to detection in noise. Journal of Neurophysiology, 118(6), 3118–3131. 10.1152/jn.00521.2017

Cohen, Y. E., Bennur, S., Christison-Lagay, K., Gifford, A. M., & Tsunada, J. (2016). Functional organization of the ventral auditory pathway. Advances in Experimental Medicine and Biology, 894, 381–388. 10.1007/978-3-319-25474-6_40

Cohen, Y. E., Russ, B. E., Gifford, G. W., 3rd, Kiringoda, R., & MacLean, K. A. (2004). Selectivity for the spatial and nonspatial attributes of auditory stimuli in the ventrolateral prefrontal cortex. The Journal of Neuroscience: The Official Journal of the Society for Neuroscience, 24(50), 11307–11316. 10.1523/JNEUROSCI.3935-04.2004

Disney, A. A., Aoki, C., & Hawken, M. J. (2012). Cholinergic suppression of visual responses in primate V1 is mediated by GABAergic inhibition. Journal of Neurophysiology, 108(7), 1907–1923. 10.1152/jn.00188.2012

Escabí, M. A., & Schreiner, C. E. (2002). Nonlinear spectrotemporal sound analysis by neurons in the auditory midbrain. The Journal of Neuroscience: The Official Journal of the Society for Neuroscience, 22(10), 4114–4131. 10.1523/jneurosci.22-10-04114.2002

Freeman, J. A., & Nicholson, C. (1975). Experimental optimization of current source-density technique for anuran cerebellum. Journal of Neurophysiology, 38(2), 369–382. 10.1152/jn.1975.38.2.369

Frey, S., Pandya, D. N., Chakravarty, M. M., Bailey, L., Petrides, M., & Collins, D. L. (2011). An MRI based average macaque monkey stereotaxic atlas and space (MNI monkey space). NeuroImage, 55(4), 1435–1442. 10.1016/j.neuroimage.2011.01.040

Fries, P. (2015). Rhythms for cognition: Communication through coherence. Neuron, 88(1), 220–235. 10.1016/j.neuron.2015.09.034

Fritz, J. B., Elhilali, M., David, S. V., & Shamma, S. A. (2007). Auditory attention-- focusing the searchlight on sound. Current Opinion in Neurobiology, 17(4), 437–455. 10.1016/j.conb.2007.07.011

Gerbella, M., Belmalih, A., Borra, E., Rozzi, S., & Luppino, G. (2010). Cortical connections of the macaque caudal ventrolateral prefrontal areas 45A and 45B. Cerebral Cortex (New York, N.Y.: 1991), 20(1), 141–168. 10.1093/cercor/bhp087

Guo, L., Weems, J. T., Walker, W. I., Levichev, A., & Jaramillo, S. (2019). Choice-selective neurons in the auditory cortex and in its striatal target encode reward expectation. The Journal of Neuroscience: The Official Journal of the Society for Neuroscience, 39(19), 3687–3697. 10.1523/JNEUROSCI.2585-18.2019

Hackett, T. A., Stepniewska, I., & Kaas, J. H. (1999). Prefrontal connections of the parabelt auditory cortex in macaque monkeys. Brain Research, 817(1-2), 45–58. 10.1016/s0006-8993(98)01182-2

Hannemann, R., Obleser, J., & Eulitz, C. (2007). Top-down knowledge supports the retrieval of lexical information from degraded speech. Brain Research, 1153, 134–143. 10.1016/j.brainres.2007.03.069

Jensen, O., & Mazaheri, A. (2010). Shaping functional architecture by oscillatory alpha activity: gating by inhibition. Frontiers in Human Neuroscience, 4, 186. 10.3389/fnhum.2010.00186

Johnston, J. M., Cohen, Y. E., Shirley, H., Tsunada, J., Bennur, S., Christison-Lagay, K., & Veeder, C. L. (2016). Recent refinements to cranial implants for rhesus macaques (Macaca mulatta). Lab Animal, 45(5), 180–186. 10.1038/laban.997

Kaas, J. H., & Hackett, T. A. (1998). Subdivisions of auditory cortex and levels of processing in primates. Audiology & Neuro-Otology, 3(2-3), 73–85. 10.1159/000013783

Logothetis, N. K. (2003). The underpinnings of the BOLD functional magnetic resonance imaging signal. The Journal of Neuroscience: The Official Journal of the Society for Neuroscience, 23(10), 3963–3971. 10.1523/jneurosci.23-10-03963.2003

Logothetis, N. K., Pauls, J., Augath, M., Trinath, T., & Oeltermann, A. (2001). Neurophysiological investigation of the basis of the fMRI signal. Nature, 412(6843), 150–157. 10.1038/35084005

Mackey, C. A., O’Connell, M. N., Hackett, T. A., Schroeder, C. E., & Kajikawa, Y. (2024). Laminar organization of visual responses in core and parabelt auditory cortex. Cerebral Cortex (New York, N.Y.: 1991), 34(9). 10.1093/cercor/bhae373

Miller, E. K., Erickson, C. A., & Desimone, R. (1996). Neural mechanisms of visual working memory in prefrontal cortex of the macaque. The Journal of Neuroscience: The Official Journal of the Society for Neuroscience, 16(16), 5154–5167. 10.1523/jneurosci.16-16-05154.1996

Mitzdorf, U. (1985). Current source-density method and application in cat cerebral cortex: investigation of evoked potentials and EEG phenomena. Physiological Reviews, 65(1), 37–100. 10.1152/physrev.1985.65.1.37

Mohn, J. L., Downer, J. D., O’Connor, K. N., Johnson, J. S., & Sutter, M. L. (2021). Choice-related activity and neural encoding in primary auditory cortex and lateral belt during feature-selective attention. Journal of Neurophysiology, 125(5), 1920–1937. 10.1152/jn.00406.2020

Moran, P. A. P. (1950). A test for the serial independence of residuals. Biometrika, 37(1-2), 178–181. 10.1093/biomet/37.1-2.178

Nicholson, C., & Freeman, J. A. (1975). Theory of current source-density analysis and determination of conductivity tensor for anuran cerebellum. Journal of Neurophysiology, 38(2), 356–368. 10.1152/jn.1975.38.2.356

Oostenveld, R., Fries, P., Maris, E., & Schoffelen, J.-M. (2011). FieldTrip: Open source software for advanced analysis of MEG, EEG, and invasive electrophysiological data. Computational Intelligence and Neuroscience, 2011, 156869. 10.1155/2011/156869

Park, H., Ince, R. A. A., Schyns, P. G., Thut, G., & Gross, J. (2015). Frontal top-down signals increase coupling of auditory low-frequency oscillations to continuous speech in human listeners. Current Biology, 25(12), 1649–1653. 10.1016/j.cub.2015.04.049

Patel, P., van der Heijden, K., Bickel, S., Herrero, J. L., Mehta, A. D., & Mesgarani, N. (2022). Interaction of bottom-up and top-down neural mechanisms in spatial multi-talker speech perception. Current Biology, 32(18), 3971–3986.e4. 10.1016/j.cub.2022.07.047

Plakke, B., & Romanski, L. M. (2014). Auditory connections and functions of prefrontal cortex. Frontiers in Neuroscience, 8, 199. 10.3389/fnins.2014.00199

Rauschecker, J. P., & Afsahi, R. K. (2023). Anatomy of the auditory cortex then and now. The Journal of Comparative Neurology, 531(18), 1883–1892. 10.1002/cne.25560

Rauschecker, J. P., & Tian, B. (2000). Mechanisms and streams for processing of “what” and “where” in auditory cortex. Proceedings of the National Academy of Sciences of the United States of America, 97(22), 11800–11806. 10.1073/pnas.97.22.11800

Reser, D. H., Fishman, Y. I., Arezzo, J. C., & Steinschneider, M. (2000). Binaural interactions in primary auditory cortex of the awake macaque. Cerebral Cortex (New York, N.Y.: 1991), 10(6), 574–584. 10.1093/cercor/10.6.574

Riddle, J., Scimeca, J. M., Cellier, D., Dhanani, S., & D’Esposito, M. (2020). Causal evidence for a role of theta and alpha oscillations in the control of working memory. Current Biology: CB, 30(9), 1748–1754.e4. 10.1016/j.cub.2020.02.065

Romanski, L. M., Bates, J. F., & Goldman-Rakic, P. S. (1999). Auditory belt and parabelt projections to the prefrontal cortex in the rhesus monkey. The Journal of Comparative Neurology, 403(2), 141–157. 10.1002/(sici)1096-9861(19990111)403:2<141::aid-cne1>3.0.co;2-v

Romanski, L. M., & Goldman-Rakic, P. S. (2002). An auditory domain in primate prefrontal cortex. Nature Neuroscience, 5(1), 15–16. 10.1038/nn781

Romanski, L. M., Tian, B., Fritz, J., Mishkin, M., Goldman-Rakic, P. S., & Rauschecker, J. P. (1999). Dual streams of auditory afferents target multiple domains in the primate prefrontal cortex. Nature Neuroscience, 2(12), 1131–1136. 10.1038/16056

Siegel, M., Donner, T. H., & Engel, A. K. (2012). Spectral fingerprints of large-scale neuronal interactions. Nature Reviews. Neuroscience, 13(2), 121–134. 10.1038/nrn3137

Tardiff, N., Suriya-Arunroj, L., Cohen, Y. E., & Gold, J. I. (2022). Rule-based and stimulus-based cues bias auditory decisions via different computational and physiological mechanisms. PLoS Computational Biology, 18(10), e1010601. 10.1371/journal.pcbi.1010601

Tian, B., Reser, D., Durham, A., Kustov, A., & Rauschecker, J. P. (2001). Functional specialization in rhesus monkey auditory cortex. Science (New York, N.Y.), 292(5515), 290–293. 10.1126/science.1058911

Tsunada, J., & Cohen, Y. E. (2014). Neural mechanisms of auditory categorization: from across brain areas to within local microcircuits. Frontiers in Neuroscience, 8, 161. 10.3389/fnins.2014.00161

Tsunada, J., Cohen, Y., & Gold, J. I. (2019). Post-decision processing in primate prefrontal cortex influences subsequent choices on an auditory decision-making task. eLife, 8(e46770), e46770. 10.7554/eLife.46770

Tsunada, J., Lee, J. H., & Cohen, Y. E. (2011). Representation of speech categories in the primate auditory cortex. Journal of Neurophysiology, 105(6), 2634–2646. 10.1152/jn.00037.2011

Tsunada, J., Liu, A. S. K., Gold, J. I., & Cohen, Y. E. (2016). Causal contribution of primate auditory cortex to auditory perceptual decision-making. Nature Neuroscience, 19(1), 135–142. 10.1038/nn.4195

Van Essen, D. C., & Fellerman, D. J. (1991). Distributed hierarchical processing in the primate cerebral cortex. Cerebral Cortex Jan/Feb, 1, 1–47. 10.1093/cercor/1.1.1-a

Weisz, N., Wühle, A., Monittola, G., Demarchi, G., Frey, J., Popov, T., & Braun, C. (2014). Prestimulus oscillatory power and connectivity patterns predispose conscious somatosensory perception. Proceedings of the National Academy of Sciences of the United States of America, 111(4), E417–E425. 10.1073/pnas.1317267111

Womelsdorf, T., Vinck, M., Leung, L. S., & Everling, S. (2010). Selective theta-synchronization of choice-relevant information subserves goal-directed behavior. Frontiers in Human Neuroscience, 4, 210. 10.3389/fnhum.2010.00210

Yin, P., Mishkin, M., Sutter, M., & Fritz, J. B. (2008). Early stages of melody processing: stimulus-sequence and task-dependent neuronal activity in monkey auditory cortical fields A1 and R. Journal of Neurophysiology, 100(6), 3009–3029. 10.1152/jn.00828.2007

Zekveld, A. A., Heslenfeld, D. J., Festen, J. M., & Schoonhoven, R. (2006). Top-down and bottom-up processes in speech comprehension. NeuroImage, 32(4), 1826–1836. 10.1016/j.neuroimage.2006.04.199

