## Supplementary material for "Different Sources of Expectations Differentially Modulate Communication Between Auditory and Prefrontal Cortices During Auditory Decision-Making": 2026_07_20_Area_Communication_Paper_Supplemental_Materials

**Table 1: Summary of one-sample Wilcoxon signed-rank tests for choice performance**. Distribution of session-wise Spearman's rank correlation (ρ) values testing the null hypothesis that there is not a monotonic relationship between SNR values and the proportion of high-frequency choices.

| **Monkey** | **Cue Type** | **N** | **Median ρ** | **25th Quartile** | **75th Quartile** | ***p*-value** |
|---|---|---|---|---|---|---|
| C | High LED | 19 | 0.9500 | 0.9133 | 0.9667 | $6.86 \times 10^{-05}$ |
| C | Neutral LED | 19 | 0.9500 | 0.9330 | 0.9667 | $6.82 \times 10^{-05}$ |
| C | Low LED | 19 | 0.9833 | 0.9333 | 0.9833 | $6.45 \times 10^{-05}$ |
| C | High Pretone | 19 | 0.9289 | 0.8833 | 0.9833 | $6.91 \times 10^{-05}$ |
| C | Low Pretone | 19 | 0.9456 | 0.9197 | 0.9667 | $6.90 \times 10^{-05}$ |
| M | High LED | 15 | 0.9833 | 0.9510 | 0.9927 | $3.05 \times 10^{-05}$ |
| M | Neutral LED | 15 | 0.9791 | 0.9500 | 0.9958 | $3.05 \times 10^{-05}$ |
| M | Low LED | 15 | 0.9667 | 0.9667 | 1.0000 | $3.05 \times 10^{-05}$ |
| M | High Pretone | 15 | 0.9791 | 0.9417 | 0.9959 | $3.05 \times 10^{-05}$ |
| M | Low Pretone | 15 | 0.9833 | 0.9667 | 0.9833 | $3.05 \times 10^{-05}$ |

**Table 2: Summary of one-sample Wilcoxon signed-rank tests for RT**. Distribution of session-wise Spearman's rank correlation (ρ) values testing the null hypothesis that there is not a monotonic relationship between |SNR| and RT.

| Monkey | Cue Type | N | Median ρ | 25th Quartile | 75th Quartile | p-value |
|---|---|---|---|---|---|---|
| C | High LED | 19 | -0.4000 | -0.6000 | 0.0000 | 0.0072 |
| C | Neutral LED | 19 | -0.9000 | -0.9000 | -0.4000 | $1.03 \times 10^{-04}$ |
| C | Low LED | 19 | -0.9000 | -0.9750 | -0.6000 | $6.66 \times 10^{-05}$ |
| C | High Pretone | 19 | -0.1000 | -0.6500 | 0.0000 | 0.0315 |
| C | Low Pretone | 19 | -0.3000 | -0.6750 | 0.0000 | 0.0055 |
| M | High LED | 15 | -0.8000 | -0.9000 | -0.7250 | $3.05 \times 10^{-05}$ |
| M | Neutral LED | 15 | -0.9000 | -1.000 | -0.9000 | $3.05 \times 10^{-05}$ |
| M | Low LED | 15 | -0.9000 | -1.000 | -0.8000 | $3.05 \times 10^{-05}$ |
| M | High Pretone | 15 | -0.3000 | -0.4750 | 0.1500 | 0.0754 |
| M | Low Pretone | 15 | -0.1000 | -0.4750 | 0.4000 | 0.4870 |

**Table 3: Model-fit (GLMM; binomial-logit) regressor values for the influence of the cue on choice behavior**. For informative trials, informative (blue/green) LED lights were compared to neutral (yellow) LEDs. The odds ratio quantifies the change in the odds of making a "high frequency" choice relative to the yellow neutral LED. An odds ratio > 1 indicates increased odds of a "high-frequency" choice, whereas a value < 1 indicates decreased odds of a "high-frequency" choice. The HHH and LLL pretones were compared directly (with LLL as the reference level), and the corresponding odds ratio quantifies the change in the odds of a "high-frequency" choice following HHH relative to LLL.

| **Model** | **N** | **Monkey C**<br>β (95% CI) | ***p*** | **Odds ratio (95% CI)** |
|---|---|---|---|---|
| Blue LED vs. Yellow LED | 55562 | 0.769 (0.726, 0.813) | $p<10^{-06}$ | 2.16 (2.07, 2.26) |
| Green LED vs. Yellow LED | 55562 | -0.563 (-0.607, -0.519) | $p<10^{-06}$ | 0.57 (0.55, 0.59) |
| Pretone: HHH vs. LLL | 28904 | 0.221 (0.173, 0.269) | $p<10^{-06}$ | 1.25 (1.19, 1.31) |

| **Model** | **N** | **Monkey M**<br>β (95% CI) | ***p*** | **Odds ratio (95% CI)** |
|---|---|---|---|---|
| Blue LED vs. Yellow LED | 70275 | 0.887 (0.848, 0.927) | $p<10^{-06}$ | 2.43 (2.34, 2.53) |
| Green LED vs. Yellow LED | 70275 | -0.792 (-0.831, -0.760) | $p<10^{-06}$ | 0.45 (0.44, 0.47) |
| Pretone: HHH vs. LLL | 42749 | 0.131 (0.092, 0.171) | $p<10^{-06}$ | 1.14 (1.10, 1.18) |

**Table 4: Model-fit (LMM) regressor values for the influence of the cue on response time**. For informative trials, we compared the difference in response time (ΔRT) during congruent/incongruent trials relative to those during uninformative trials. For uninformative trials, ΔRT reflects the difference between congruent and incongruent trials.

| **Model** | **N** | **Monkey C** ΔRT ms (95% CI) | ***p*** |
|---|---|---|---|
| Congruent LED vs. Neutral | 13604 | -43.4 (-52.5, -34.3) | $p<10^{-06}$ |
| Incongruent LED vs. Neutral LED | 13604 | +32.6 (20.4, 44.8) | $p<10^{-06}$ |
| Incongruent Pretone vs. Congruent Pretone | 42729 | +35.2 (23.0, 47.4) | $p<10^{-06}$ |

| **Model** | **N** | **Monkey M** ΔRT ms (95% CI) | ***p*** |
|---|---|---|---|
| Congruent LED vs. Neutral | 14265 | -34.5 (-40.0, -29.0) | $p<10^{-06}$ |
| Incongruent LED vs. Neutral LED | 14265 | +43.2 (35.8, 50.5) | $p<10^{-06}$ |
| Incongruent Pretone vs. Congruent Pretone | 6907 | +35.2 (27.6, 42.8) | $p<10^{-06}$ |

**Table 5a: Summary of response-index (RI) values**.

| Monkey | Cue Type | Area | N | Median RI | 25th Quartile | 75th Quartile |
|---|---|---|---|---|---|---|
| C | Informative | AC | 19 | 0.19390 | 0.11820 | 0.24560 |
| C | Informative | vlPFC | 19 | 0.05599 | 0.04175 | 0.06954 |
| C | Uninformative | AC | 19 | 0.03807 | 0.02231 | 0.07431 |
| C | Uninformative | vlPFC | 19 | 0.04572 | 0.03598 | 0.05893 |
| M | Informative | AC | 15 | 0.14040 | 0.12120 | 0.16840 |
| M | Informative | vlPFC | 15 | 0.06076 | 0.04899 | 0.07347 |
| M | Uninformative | AC | 15 | 0.03533 | 0.00015 | 0.13200 |
| M | Uninformative | vlPFC | 15 | 0.03416 | 0.03061 | 0.05156 |

**Table 5b: Summary of pairwise Wilcoxon signed-rank tests for RI values**. We tested the null hypothesis that RI values were the same during informative and uninformative trials as a function of cortical area (AC or vlPFC).

| Monkey | Area | N | $p$ | $z$ |
|---|---|---|---|---|
| C | AC | 19 | 0.00085 | -3.7023 |
| C | vlPFC | 19 | 0.03600 | -2.0926 |
| M | AC | 15 | 0.0076 | -2.6694 |
| M | vlPFC | 15 | 0.017 | -2.3854 |

**Table 6a: Summary of pairwise Wilcoxon signed-rank tests for high-SNR, informative trials**. Within each communication direction and congruency condition, we tested the null hypothesis that correct and incorrect trials exhibited the same median PSI values.

| Monkey | Direction | Congruency | N | Cue Type | z | *p* |
|---|---|---|---|---|---|---|
| C | Top-Down | Congruent | 2087 | Informative | 24.295 | $p<10^{-06}$ |
| C | Bottom-Up | Congruent | 2065 | Informative | 20.179 | $p<10^{-06}$ |
| M | Top-Down | Congruent | 1590 | Informative | 26.249 | $p<10^{-06}$ |
| M | Bottom-Up | Congruent | 1770 | Informative | 22.878 | $p<10^{-06}$ |
| C | Top-Down | Incongruent | 1849 | Informative | 7.950 | $p<10^{-06}$ |
| C | Bottom-Up | Incongruent | 2171 | Informative | 8.355 | $p<10^{-06}$ |
| M | Top-Down | Incongruent | 1643 | Informative | 7.792 | $p<10^{-06}$ |
| M | Bottom-Up | Incongruent | 1671 | Informative | 6.116 | $p<10^{-06}$ |

**Table 6b: Summary of pairwise Wilcoxon signed-rank tests for high-SNR, uninformative trials**. Within each communication direction and congruency condition, we tested the null hypothesis that correct and incorrect trials exhibited the same median PSI values.

| **Monkey** | **Direction** | **Congruency** | **N** | **Cue Type** | **z** | ***p*** |
|---|---|---|---|---|---|---|
| C | Top-Down | Congruent | 1670 | Uninformative | 19.663 | $p<10^{-06}$ |
| C | Bottom-Up | Congruent | 2186 | Uninformative | 21.788 | $p<10^{-06}$ |
| M | Top-Down | Congruent | 1578 | Uninformative | 20.090 | $p<10^{-06}$ |
| M | Bottom-Up | Congruent | 1462 | Uninformative | 15.642 | $p<10^{-06}$ |
| C | Top-Down | Incongruent | 2189 | Uninformative | 15.156 | $p<10^{-06}$ |
| C | Bottom-Up | Incongruent | 1664 | Uninformative | 17.019 | $p<10^{-06}$ |
| M | Top-Down | Incongruent | 1599 | Uninformative | 15.008 | $p<10^{-06}$ |
| M | Bottom-Up | Incongruent | 1419 | Uninformative | 13.143 | $p<10^{-06}$ |

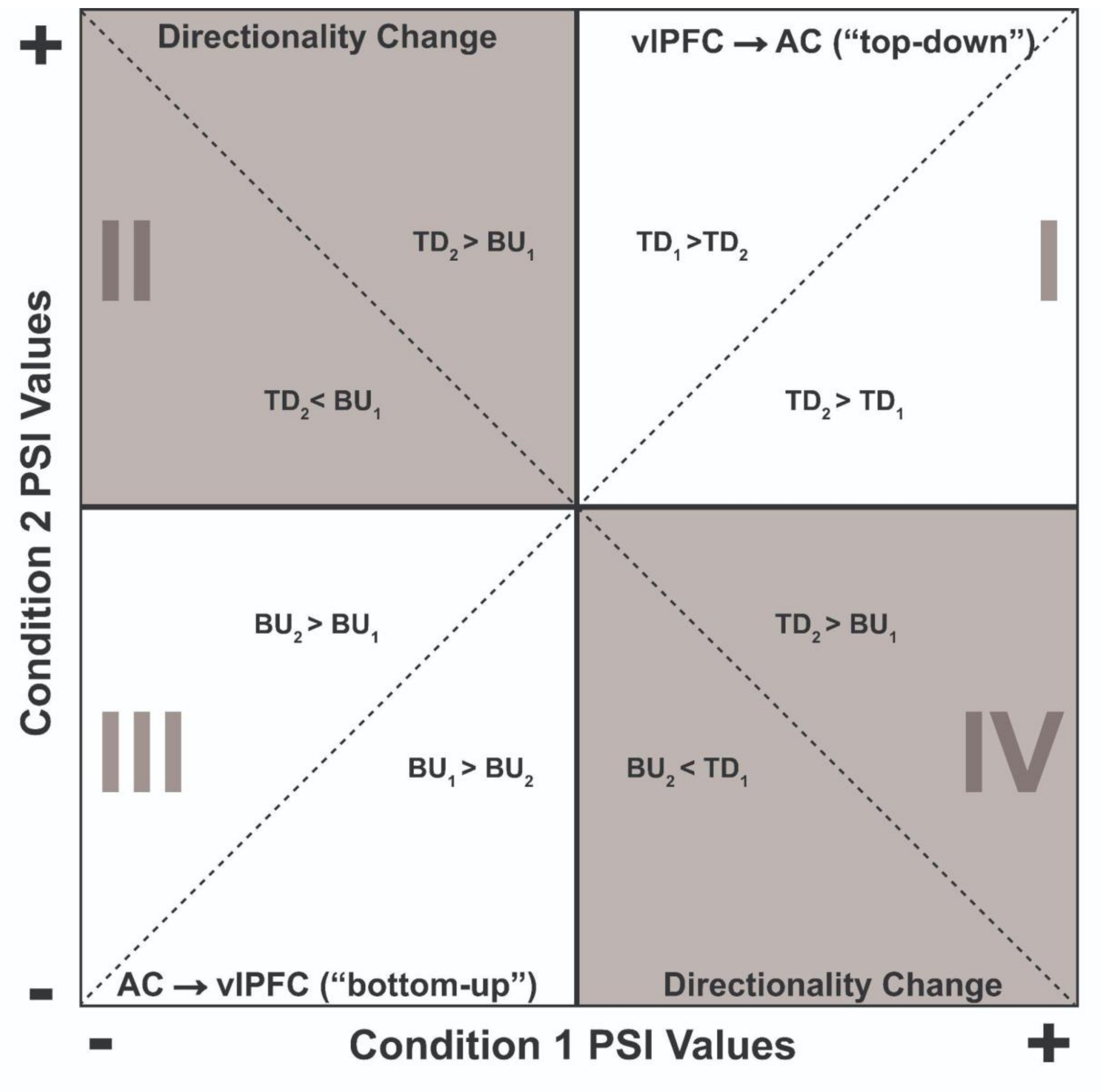


**Figure S1: Cartoon schematic of the relationship between the sign of phase slope index (PSI) and experimental condition.** Negative PSI values indicate information flow from AC→vlPFC, whereas positive PSI values indicate information flow from vlPFC→AC. PSI magnitude indicates the strength of the AC-vlPFC connection. The position of a PSI value when plotted on a cartesian plane conveys information about the directionality of information flow across different experimental conditions. Recording site pairs in quadrants I and III preserve their directionality across conditions (remaining top-down or bottom-up, respectively), whereas pairs in quadrants II and IV exhibit reversals in directionality (i.e., switching between top-down and bottom-up states). The inequalities in the figure indicate how PSI values change with respect to condition 1 and condition 2. TD: top-down. BU: bottom-up. The subscripts indicate condition 1 or condition 2.

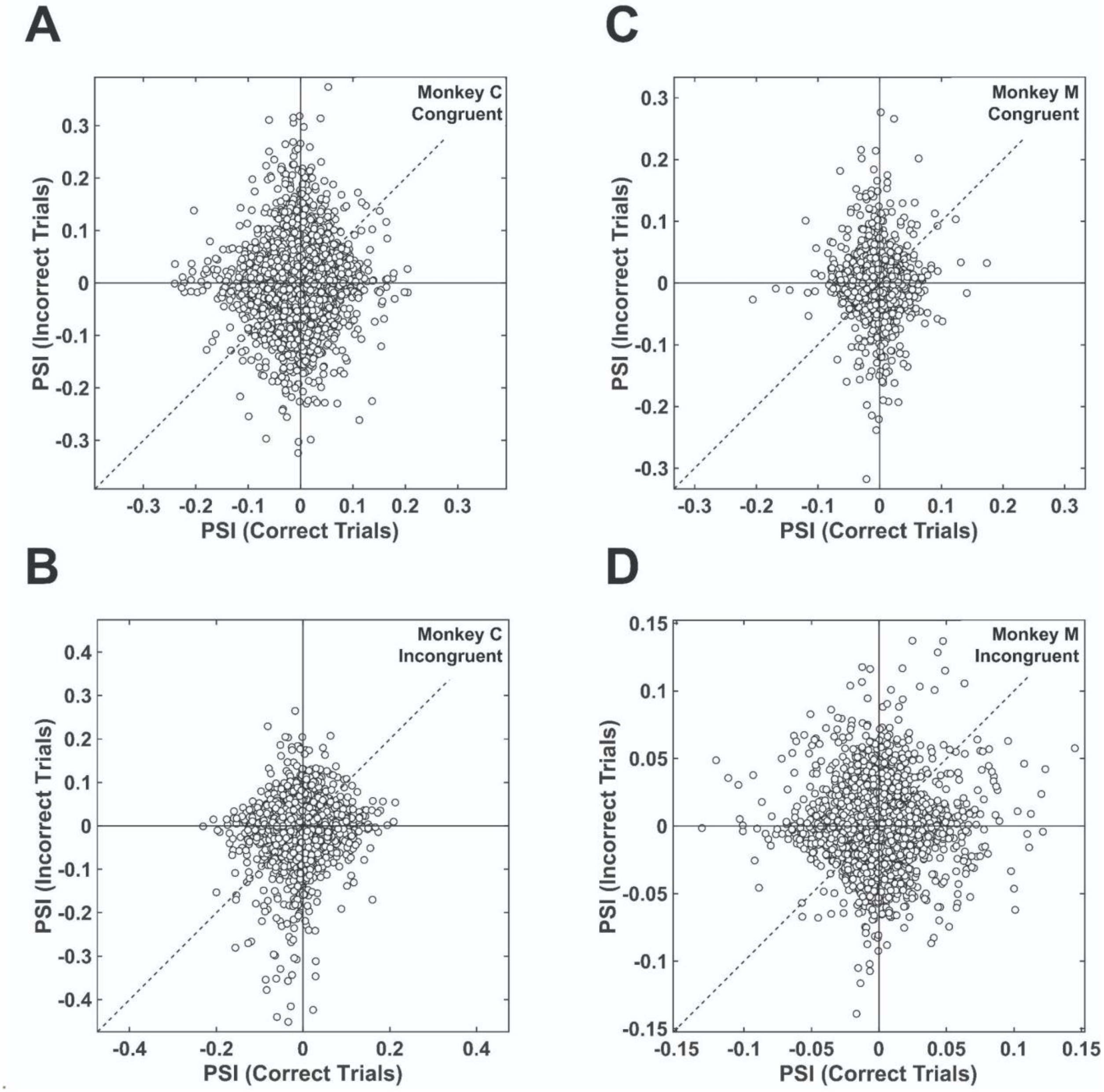


**Figure S2: During high-SNR informative trials, PSI is not modulated by congruency.** Scatter plots of PSI values during correct (x-axis) and incorrect (y-axis) trials for Monkey C (**A, B**) and Monkey M (**C, D**). Data in the **top** row are from congruent trials, whereas data in the **bottom** row are from incongruent trials. Each plot represents all the recording-site pairs from monkey C (N = 7600) and monkey M (N = 6000) pooled across sessions (Monkey C, 19 sessions; Monkey M, 15 sessions).

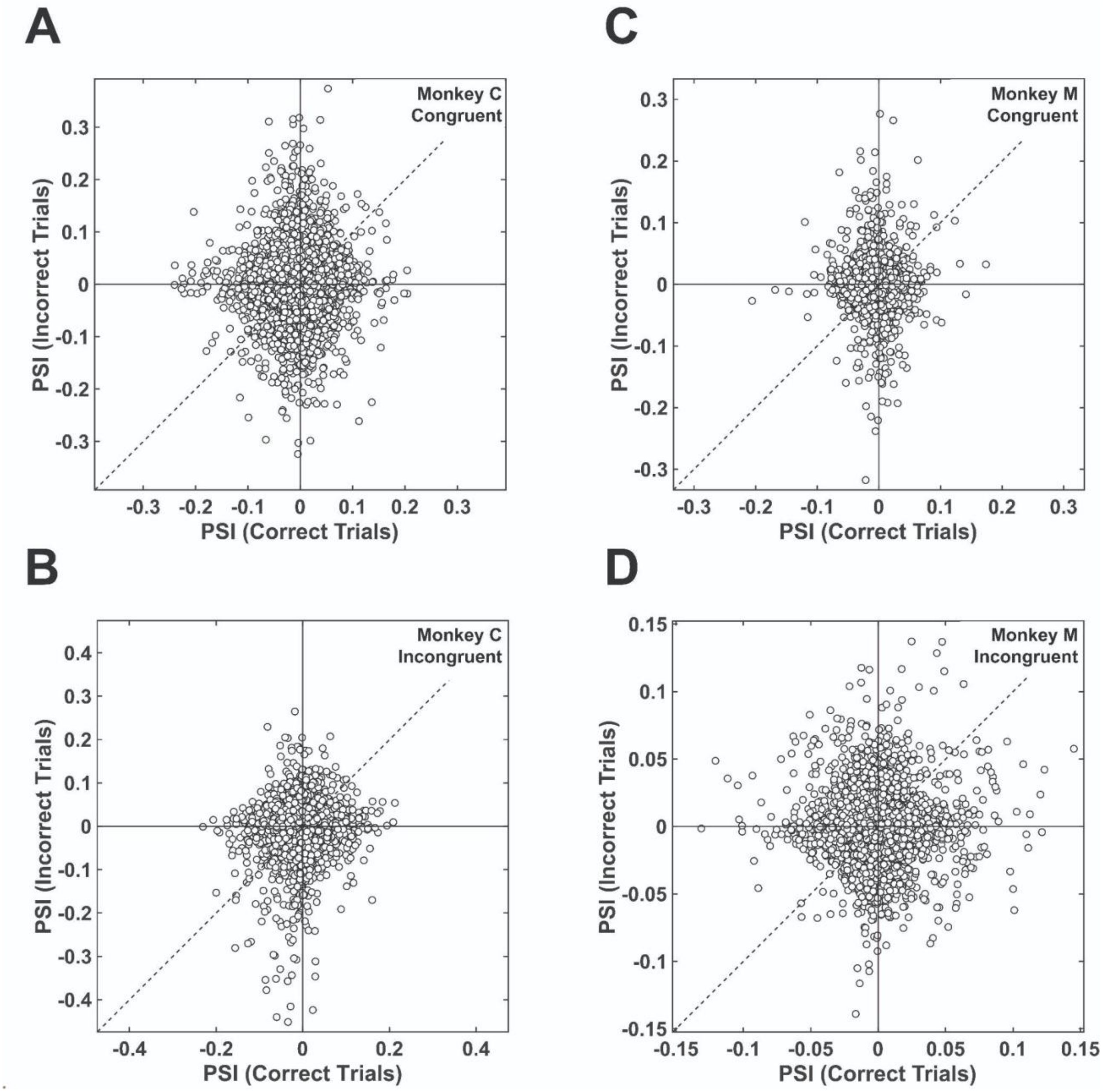


**Figure S3: During high-SNR uninformative trials, PSI is not modulated by congruency.** Scatter plots of PSI values during correct (x-axis) and incorrect (y-axis) trials for Monkey C (**A, B**) and Monkey M (**C, D**). Data in the **top** row are from congruent trials, whereas data in the **bottom** row are from incongruent trials. Each plot represents all the recording-site pairs from monkey C (N = 7600) and monkey M (N = 6000) pooled across sessions (Monkey C, 19 sessions; Monkey M, 15 sessions).

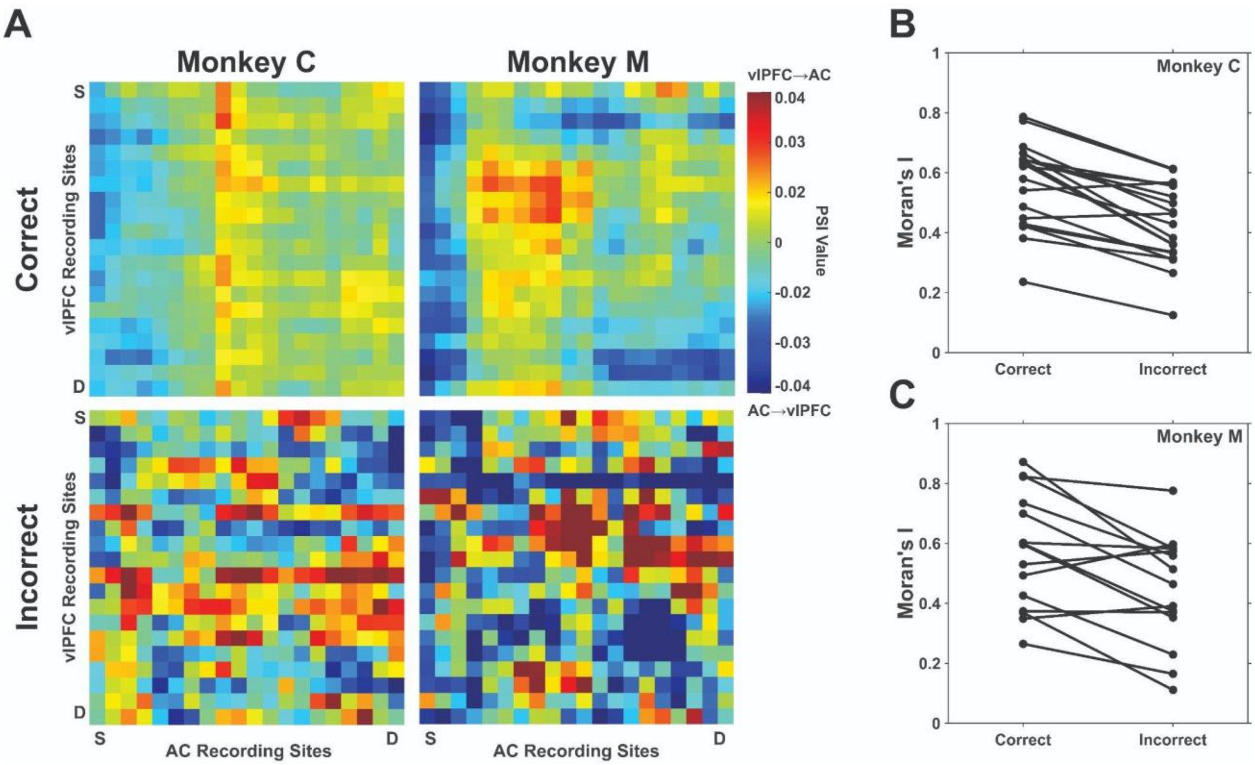


**Figure S4: During high-SNR, congruent informative trials, spatial organization of PSI values are modulated by outcome.** (**A**) Across-session mean PSI values (Monkey C, N = 19, Monkey M, N = 15) during the target-tone period as a function of each recording-site pair. These pairs are organized relative to each AC and vlPFC probe's position in the cortex from superficial (S) to deep (D). Positive PSI values are indicated by warm colors and indicate vlPFC→AC interactions, whereas negative PSI values are indicated by cool colors and indicate AC→vlPFC interactions. The spatial organization of PSI values during correct trials are plotted in the **top** row, whereas the spatial organization of PSI values during incorrect trials are plotted in the **bottom** row. (**B, C**). Session-by-session Moran's I values for correct and error trials. Moran's I values from the same session are connected by a line. A Moran's I value of -1 indicates that the spatial organization of the data is dispersed, 0 indicates randomness, and 1 indicates clustered.
